# Peptide vaccine candidate mimics the heterogeneity of natural SARS-CoV-2 immunity in convalescent humans and induces broad T cell responses in mice models

**DOI:** 10.1101/2020.10.16.339937

**Authors:** Eszter Somogyi, Zsolt Csiszovszki, Levente Molnár, Orsolya Lőrincz, József Tóth, Sofie Pattijn, Jana Schockaert, Aurélie Mazy, István Miklós, Katalin Pántya, Péter Páles, Enikő R. Tőke

## Abstract

We developed a global peptide vaccine against SARS-CoV-2 that addresses the dual challenges of heterogeneity in the immune responses of different individuals and potential heterogeneity of the infecting virus. PolyPEPI-SCoV-2 is a polypeptide vaccine containing nine 30-mer peptides derived from all four major structural proteins of the SARS-CoV-2. Vaccine peptides were selected based on their frequency as HLA class I and class II personal epitopes (PEPIs) restricted to multiple autologous HLA alleles of individuals in an *in silico* cohort of 433 subjects of different ethnicities. PolyPEPI-SCoV-2 vaccine administered with Montanide ISA 51VG adjuvant generated robust, Th1-biased CD8^+^ and CD4^+^ T cell responses against all four structural proteins of the virus, as well as binding antibodies upon subcutaneous injection into BALB/c and CD34^+^ transgenic mice. In addition, PolyPEPI-SCoV-2-specific, polyfunctional CD8^+^ and CD4^+^ T cells were detected *ex vivo* in each of the 17 asymptomatic/mild COVID-19 convalescents’ blood investigated, 1–5 months after symptom onset. The PolyPEPI-SCoV-2-specific T cell repertoire used for recovery from COVID-19 was extremely diverse: donors had an average of seven different peptide-specific T cells, against the SARS-CoV-2 proteins; 87% of donors had multiple targets against at least three SARS-CoV-2 proteins and 53% against all four. In addition, PEPIs determined based on the complete HLA class I genotype of the convalescent donors were validated, with 84% accuracy, to predict PEPI-specific CD8^+^ T cell responses measured for the individuals. Extrapolation of the above findings to a US bone marrow donor cohort of 16,000 HLA-genotyped individuals with 16 different ethnicities (n=1,000 each ethnic group) suggest that PolyPEPI-SCoV-2 vaccination in a general population will likely elicit broad, multi-antigenic CD8^+^ and CD4^+^ T cell responses in 98% of individuals, independent of ethnicity, including Black, Asian, and Minority Ethnic (BAME) cohorts.

## Introduction

The pandemic caused by the novel coronavirus SARS-CoV-2 is still evolving after its outbreak in December 2019. According to World Health Organization (WHO), at least two-thirds of the vaccine candidates under clinical development are designed to generate primarily neutralizing antibodies against the viral Spike (S) protein (*1*), but lessons learned from the SARS and MERS epidemic as well as COVID-19 convalescents indicate potential challenges for this vaccine design strategy.(*2, 3*) Potential issues are dual: waning antibody levels and inefficient T cell response generation against only the Spike protein.

Patients infected with the previous SARS-CoV endemic in 2003 and MERS endemic in 2012 often had transient (detected only for up to 3-6 years) humoral immunity.(*4*)(*5*) Even more, antibodies generated by a low-risk experimental infection with a common cold coronavirus declined within 1 year and did not protect against re-challenge.(*6, 7*) Similarly, with SARS-CoV-2, immune responses associated with the natural course of SARS-CoV-2 viral infection suggest that anti-Spike IgG antibody responses are usually weak (except for the fortunately less frequent severe cases) and their durability lasts for up to 3 months, in most cases, or decline by up to 70% within this time period.(*8*) In addition, 2–9% of individuals do not seroconvert even 2 months after infection with SARS-CoV-2,(*9*) suggesting that individuals reached immunity using another arm of the adaptive immune system, T cells. Indeed, it could be concluded that virtually all subjects with a history of SARS-CoV-2 infection mount T cell responses against the virus, including seronegatives and subjects with severe disease.(*2, 10–12*) T cell responses are diverse, i.e., directed against the whole antigenic repertoire of the virus, and less dominated by S protein. Specifically, several studies reported that despite being the largest structural protein, S-specific T cell responses accounted for only 25–27% of the totality of CD4^+^ and CD8^+^ T cells elicited by the natural infection. Furthermore, this diversity is associated with asymptomatic/mild disease as recovered patients had more CD8^+^ T cells against Membrane (M) and Nucleoprotein (N) proteins rather than S, while T cell intensity and diversity do not increase with disease severity, as demonstrated for MERS, SARS-CoV-1, and SARS-CoV-2.(*8, 11, 13, 14*) Indeed, in COVID-19 patients, low CD8^+^ T cell counts are a predictor of higher risk for death, and patients with severe disease or who died had exhausted T cells.(*2, 15*) It was proposed that detectable virus-specific CD8^+^ T cell responses at earlier times after infection contribute to lower viral load and therefore lower antibody levels, explaining why these patients have more favorable outcomes.(*14*) In support of this, it was recently reported that mapping of SARS-CoV-2-specific T cell receptors was possible soon after viral exposure and prior to any antibody detection.(*16*) These observations also suggest that achieving elevated numbers of diverse virus-specific memory T cells prior to infection (by vaccination) may contribute to virus- and viral reservoir elimination in the early-stage of SARS-CoV-2 infection. These expectations are supported by animal challenge studies demonstrating that reactivated T cells provided protection from lethal dose infection with SARS-CoV.(*11, 17*) Remarkably, memory T cells against the N protein of SARS-CoV were reported for 23/23 patients tested 17 years after their recovery from SARS.(*11*) Other reports also supported the durability of memory T cells elicited by coronavirus infections.(*4, 18*)

Therefore, vaccine candidates under clinical development aiming to generate T cell responses against the viral S protein will likely only generate a fraction of the convalescent’s immune responses, and therefore less likely induce robust memory T cell responses. Vaccine technologies using whole viruses or multiple large proteins could theoretically solve the issue related to lack of diversity. However, these have the limitation of inclusion of unnecessary antigenic load that not only contributes little to the protective immune response, but complicates the situation by inducing allergenic and/or reactogenic responses.(*19–24*) Indeed, for SARS, it was suggested that candidate coronavirus vaccines that limit the inclusion of whole viral proteins may have more beneficial safety profiles.(*25, 26*) Similarly, replication-deficient viral constructs encoding target antigens could trigger unspecific immune responses against the viral vector, especially with repeated doses.(*27*)

Peptide vaccines are an attractive alternative subunit vaccine strategy that relies on use of short peptide fragments and epitopes that are capable of inducing positive, desirable T cell- and B cell-mediated immune responses. The core problem that afflicts peptide vaccine design, however, is that each human has a uniquely endowed immune response profile. Indeed, for SARS-CoV-2, the disease course varies according to the genetic diversity represented by different ethnicities, especially Black, Asian and Minority Ethnic (BAME) groups; however, the reason for this is not yet well understood.(*28, 29*) Genetic diversity could be captured by genetic variance in human leukocyte antigen (HLA) alleles, which are critical components of the viral antigen (epitope) presentation pathway that triggers the cytotoxic T cells (CTLs) capable of recognizing and killing cancer or infected cells in the body. To capture this heterogeneity in the design of a global vaccine against SARS-CoV-2, viral antigen epitope prediction based on frequent human HLA alleles has been used widely.(*30*) However, in reality, these epitope mapping studies have a low yield in terms of validated T cell responses. For example, in one study, 100 SARS-CoV-2-derived epitopes predicted for the 10 most prevalent HLA class I alleles were tested and only 12 were confirmed as dominant epitopes, i.e., recognized by >50% of COVID-19 donor CD8^+^ T cells. This is consistent with immune response rates observed in the field for several infectious disease and cancer vaccine clinical trials, as well as for the relatively low confirmation level of personalized mutational neoantigen-based epitopes.*(31–33)*

To overcome these limitations of peptide vaccine design, we developed PolyPEPI-SCoV-2 digitally using ethnically diverse *in silico* human cohort of individuals with complete HLA genotypes, instead using single HLA alleles. We selected multiple so-called Personal Epitopes (PEPIs) restricted to not only one but multiple autologous HLA alleles of each individual but that are also shared among a high proportion of subjects in an ethnically diverse population. Notably, this *in silico* human cohort together with the PEPI concept previously retrospectively predicted the immune response rates of 79 vaccine clinical trials, as well as the remarkable immunogenicity (80% CD8^+^ T cell responses against at least three out of six antigens) of our PolyPEPI1018 cancer vaccine in a clinical trial conducted in metastatic colorectal cancer patients.*(34–36)* CD8^+^ T cell responses generated by PEPIs in a personalized polypeptide mixture prepared for a patient with breast cancer proved to be long-lasting as they were detected 14 months after last vaccination against four tumor antigens.(*37*)

Consistent with the apparent long-term memory T cell formation capacity of SARS-CoV-2 during the natural course of infection, we designed a polypeptide vaccine to: (1) induce robust and broad immune responses in each subject by targeting all four structural proteins of SARS-CoV-2; (2) address and overcome the potential virus evolution effect by ensuring multiple immunogenic target in each patient; and (3) address different sensitivities of human ethnicities by Personal Epitope coverage of the peptides. Here, we report the design and preclinical characterization of our vaccine candidate against COVID-19. Immunogenicity and safety were confirmed in two mouse models, resulting in the induction of robust CD4^+^ and CD8^+^ T cell responses boosted by the second dose, as well as humoral responses. In convalescent COVID-19 blood samples, vaccine-specific immune cells were detected against all peptides and in all subjects, representing important components of the SARS-CoV-2-induced immune repertoire leading to recovery from infection. Peptide vaccines are a safe and economic technology compared with traditional vaccines made of dead or attenuated viruses and recombinant proteins. Synthetic peptide manufacturing at a multi-kilogram scale is relatively inexpensive and more mature than mRNA production. Our technology enables not only identification of the antigen targets for a specific disease/pathogen but, more importantly, computational determination of the antigens that immune systems of individuals in large cohorts can respond to.

## Materials and Methods

### Donors

Donors were recruited based on their clinical history of SARS-CoV-2 infection. Blood samples were collected from convalescent individuals (n=15) at an independent medical research center in The Netherlands under an approved protocol (NL57912.075.16.) or collected by PepTC Vaccines Ltd (n=2). Sera and PBMC samples from non-exposed individuals (n=5) were collected before 2018 and were provided by Nexelis-IMXP (Belgium). All donors provided written informed consent. The study was conducted in accordance with the Declaration of Helsinki. Blood samples from COVID-19 convalescent patients (n=17; 16 with asymptomatic/mild disease and one with severe disease) were obtained 17–148 days after symptom onset. Surprisingly, one positive IgM antibody response was found among the healthy donors, which was excluded from further analysis. Demographic and baseline information of the subjects are provided in Supplementary Table 1. HLA genotyping of the convalescent donor patients from The Netherlands was done by IMGM laboratories GmbH (Martinsried, Germany) using next-generation sequencing.

### Animals

#### CD34^+^ transgenic humanized mouse (Hu-mouse)

*Female* NOD/Shi-scid/IL-2Rγ null immunodeficient mice (Charles River Laboratories, France) were humanized using hematopoietic stem cells (CD34^+^) isolated from human cord blood. Only mice with a humanization rate (hCD45/total CD45) >50% were used during the study. Experiments were carried out with 20–23-week-old female mice.

#### BALB/c mouse

Experiments were carried out with 6–8-week-old female BALB/c mice (Janvier, France).

### Vaccine design

SARS-CoV-2 structural proteins (S, N, M, E) were screened and nine different 30-mer peptides were selected during a multi-step process. First, sequence diversity analysis was performed (as of 28 March 2020 in the NCBI database).(*38*) The accession IDs were as follows: **NC_045512.2**, MN938384.1, MN975262.1, MN985325.1, MN988713.1, MN994467.1, MN994468.1, MN997409.1, MN988668.1, MN988669.1, MN996527.1, MN996528.1, MN996529.1, MN996530.1, MN996531.1, MT135041.1, MT135043.1, MT027063.1, and MT027062.1. The bolded ID represents the GenBank reference sequence. Then, the translated coding sequences of the four structural protein sequences were aligned and compared using a multiple sequence alignment (Clustal Omega, EMBL-EBI, United Kingdom).(*39*) Of the 19 sequences, 15 were identical; however, single AA changes occurred in four N protein sequences: MN988713.1, N 194 S->X; MT135043.1, N 343 D->V; MT027063.1, N 194 S->L; MT027062.1, N 194 S->L. The resulting AA substitutions affected only two positions of N protein sequence (AA 194 and 343), neither of which occurred in epitopes that have been selected as targets for vaccine development. Only one (H49Y) of the 13 reported single-letter changes in the viral S protein (D614G, S943P, L5F, L8V, V367F, G476S, V483A, H49Y, Y145H/del, Q239K, A831V, D839Y/N/E, P1263L) has been involved in the PolyPEPI-SCoV-2 vaccine, but the prevalence of this variant is decreasing among later virus isolates.(*40*) Further details on peptide selection are provided in the Results section and the resulting composition of the nine selected 30-mer peptides is shown in Table 1.

**Table 1.**
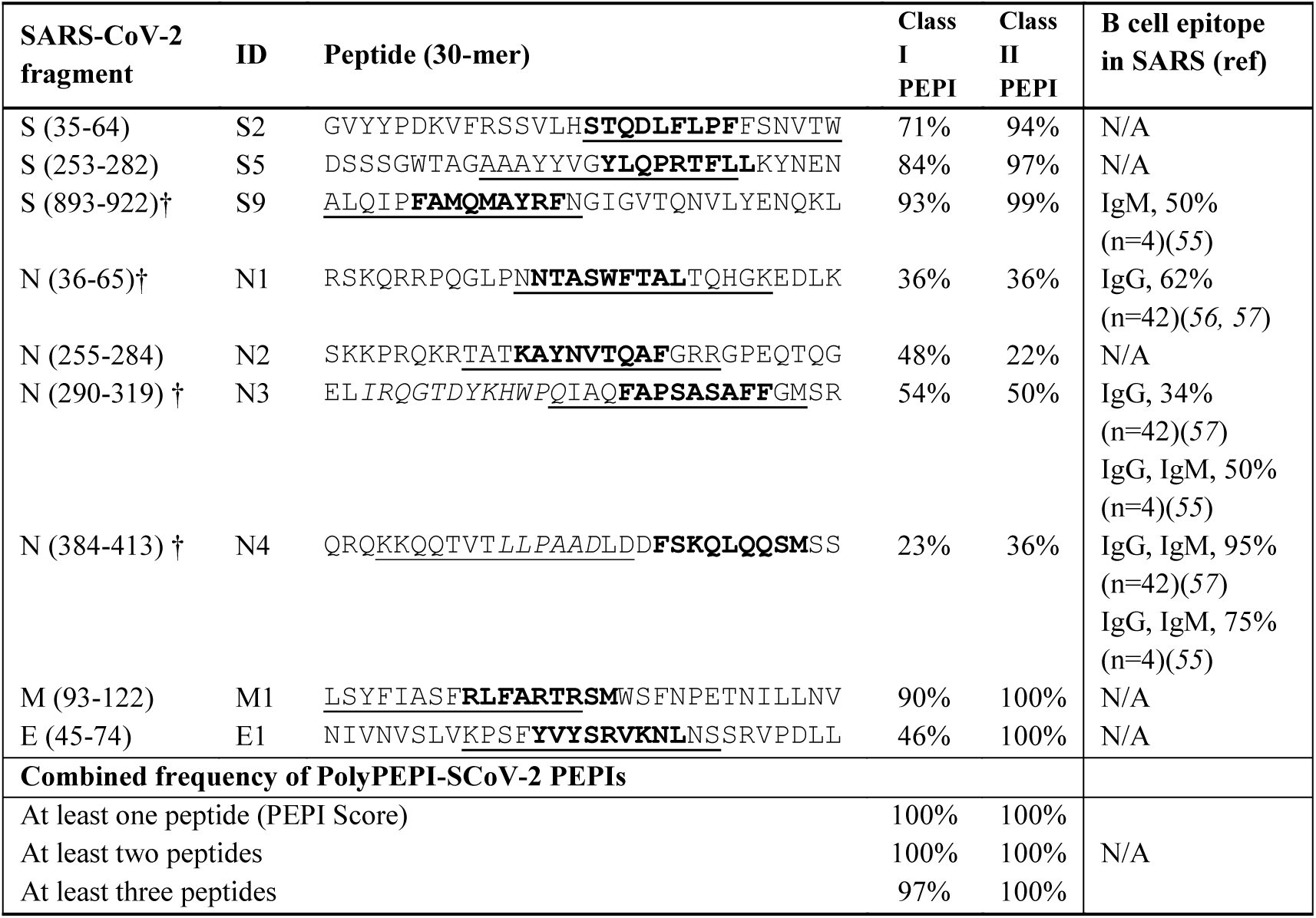
PolyPEPI-SCoV-2 peptides and comprising PEPI frequencies within the *in silico* human cohort. Bold: 9-mer HLA class I PEPI sequences within PolyPEPI-SCoV-2 comprising 30-mer peptides; underlined: 15-mer HLA class II PEPI sequences. Percentages show the proportion of individuals from the model population with at least one HLA class I (CD8^+^ T cell specific) PEPI or at least one HLA class II (CD4^+^ T cell specific) PEPI. Peptides labeled † contain experimentally confirmed B cell epitopes with antibody (Ig) responses found in convalescent patients with SARS. N/A: data not available.

### Cross-reactivity with human coronavirus strains

The sequence of PolyPEPI-SCoV-2 vaccine was compared with that of SARS-CoV, MERS-CoV, and common (seasonal) human coronavirus strains to reveal possible cross-reactive regions. According to Centers for Disease Control and Prevention (CDC), common coronaviral infections in the human population are caused by four coronavirus groups: alpha coronavirus 229E and NL63, and beta coronavirus OC43 and HKU1.(*41*) Pairwise alignment of the structural proteins was also performed using Uniprot (*42*) with the following reference sequence IDs: 229E: P15423 (S), P15130 (N), P19741 (E), P15422 (M); NL63: Q6Q1S2 (S), Q6Q1R8 (N), Q6Q1S0 (E), Q6Q1R9 (M); OC43: P36334 (S), P33469 (N), Q04854 (E), Q01455 (M); HKU1 (Isolate N1): Q5MQD0 (S), Q5MQC6 (N), Q5MQC8 (E), Q5MQC7 (M). In addition, the coronavirus strains were aligned with the nine 30-mer peptides comprising the PolyPEPI-SCoV-2 vaccine. For the minimum requirement of an epitope, eight AA-long regions were screened (sliding window) as regions responsible for potential cross-reactivity. In addition, shorter (and longer) length matching consecutive peptide fragments were recorded and reported.

### In *silico* human cohorts

The Model Population is a cohort of 433 individuals representing several ethnic groups worldwide for whom complete HLA class I genotypes were available (2 × HLA-A, 2 × HLA-B, 2 × HLA-C). The Model Population was assembled from 90 Yoruban African (YRI), 90 European (CEU), 45 Chinese (CHB), 45 Japanese (JPT), 67 subjects with mixed ethnicity (US, Canada, Australia, New Zealand), and 96 subjects from an HIV database (MIX).(*43–46*) HLA genotypes were determined using PCR techniques, Affymetrix 6.0 and Illumina 1.0 Million SNP mass arrays, and high-resolution HLA typing of the six HLA genes by Reference Strand-mediated Conformational Analysis (RSCA) or sequencing-based typing (SBT).(*47–49*) Characterization of the Model Population was reported previously.(*34*)

A second cohort of 356 individuals with characterized HLA class II genotypes (2 × HLA-DRB, 2 × HLA-DP, and 2 × HLA-DQ) at four-digit allele resolution was obtained from the dbMHC database(*50*), an online available repository operated by the National Center for Biotechnology Information (NCBI). HLA genotyping was performed by SBT.

*Large, US cohort (n=16,000)*

The database comprising anonymized HLA-genotype data from 16,000 individuals was created by obtaining 1,000 donors from each of 16 ethnic groups (500 male and 500 female) from the National Marrow Donor Program (NMDP).(*51*) The 16 ethnic groups were: African, African American, Asian Pacific Islander, Filipino, Black Caribbean, Caucasian, Chinese, Hispanic, Japanese, Korean, Native American Indian, South Asian, Vietnamese, US, Mideast/North coast of Africa, Hawaiian, and other Pacific Islander. The ethnic groups represented in this large US cohort covers the composition of the global population but they were not weighted for their global representativeness (we intentionally used n=1,000 subjects for each ethnicity).(*52*) HLA genotyping was performed by NMDP recruitment labs using sequence-specific oligonucleotide (SSO) and sequence specific primer (SSP) methods with an average “typing resolution score” >0.7.(*53*)

### Peptides and PolyPEPI-SCoV-2 vaccine preparation

The 9-mer (s2, s5, s9, n1, n2, n3, n4, e1, m1) and 30-mer (S2, S5, S7, N1, N2, N3, N4, E1, M1) peptides were manufactured by Intavis Peptide Services GmbH&Co. KG (Tübingen, Germany) and PEPScan (Lelystad, The Netherlands) using solid-phase peptide synthesis. Amino acid sequences are provided in Table 1 for both 9-mer test peptides (Table 1, bold) and the 30-mer vaccine peptides. Research grade PolyPEPI-SCoV-2 vaccine for the animal study was prepared by dissolving equal masses of the nine 30-mer peptides in DMSO to achieve at a concentration of 1 mg/mL and then diluted with purified water to a final concentration of 0.2 mg/mL and stored frozen until use. Ready-to-inject vaccine preparations were prepared by emulsifying equal volumes of thawed peptide mix solution and Montanide ISA 51 VG adjuvant (Seppic, Paris, France) following the standard two-syringe protocol provided by the manufacturer.

### Epitope prediction and analysis

Prediction of ≥3HLA class I allele binding epitopes (PEPIs) for each individual was performed using an Immune Epitope Database (IEDB)-based epitope prediction method. The antigens were scanned with overlapping 9-mer and 15-mer peptides to identify peptides that bind to the subject’s HLA class I alleles. Selection parameters were validated with an in-house set of 427 HLA-epitope pairs that had been characterized experimentally by using direct binding assays (327 binding and 100 non-binding HLA-epitope pairs). A specificity and sensitivity of 93% was achieved for the prediction of true HLA allele-epitope pairs. HLA class II epitope predictions were performed by NetMHCpan (2.4) prediction algorithm for overlapping 15-mer peptides. ≥4 HLA class II binding epitopes per individual are defined as HLA class II PEPI.

### Preclinical animal study design

Thirty-six Hu-mice and 36 BALB/c mice received PolyPEPI-SCoV-2 vaccine (0.66 mg/kg/peptide in 200 µL solution; n=18) or 20% DMSO/water emulsified in Montanide ISA 51 VG adjuvant (200 µL vehicle; n=18) administered subcutaneously on days 0 and 14; the follow up period ended on day 28. Samples from days 14, 21, and 28 were analyzed (n=6 per cohort). The studies were performed at the Transcure Bioservices facility (Archamps, France). The mice were monitored daily for unexpected signs of distress. Complete clinical scoring was performed weekly by monitoring coat (score 0–2), movement (0–3), activity (0– 3), paleness (0–2), and bodyweight (0–3); a normal condition was scored 0.

All procedures described in this study have been reviewed and approved by the local ethic committee (CELEAG) and validated by the French Ministry of Research. Vaccination-induced T cell responses were assessed by *ex vivo* ELISpot and intracellular cytokine staining (ICS) assays of mice splenocytes (detailed below). Antibody responses were investigated by the measurement of total IgG in plasma samples (detailed below).

### ELISpot/FluoroSpot assays

*Ex vivo* ELISpot assays for animal studies were performed as follows. IFN-γ-producing T cells were identified using 2 × 10^5^ splenocytes stimulated for 20 h/peptide (10 μg/ml, final concentration). Splenocytes were treated with 9-mer peptides (a pool of four N-specific peptides, N-pool [n1, n2, n3, n4], a pool of three S-specific peptides, S-pool [s2, s5, s9], an E protein-derived peptide, e1 or a M protein-derived peptide, m1) or with 30-mer peptides pooled the same way as 9-mers (N-pool comprising peptides N1, N2, N3, and N4), S-pool comprising peptides S2, S5 and S9, and individual peptides E1 and M1. ELISpot assays were performed using Human IFN-γ ELISpot PRO kit (ALP; ref 3321-4APT-2) from mabTech for Hu-mice cohorts and Mouse IFN-γ ELISpot PRO kit (ALP; ref 3321-4APT-10) from mabTech for BALB/c mice cohorts, according to the manufacturer’s instructions. Unstimulated (DMSO) assay control background spot forming unit (SFU) was subtracted from each data point and then the delta SFU (dSFU) was calculated. PMA/Ionomycin (Invitrogen) was used as a positive control.

*Ex vivo* FluoroSpot assays for convalescent donor testing were performed by Nexelis-IMXP (Belgium) as follows: IFN-γ/IL-2 FluoroSpot plates were blocked with RPMI-10% FBS, then peptides (5 µg/mL final concentration) or peptide pools (5 µg/mL per peptide final concentration) were added to the relevant wells. PBMCs were retrieved from cryogenic storage and thawed in culture medium. Then, 200,000 PBMC cells/well were plated in triplicate (stimulation conditions) or 6-plicates (reference conditions) and incubated overnight at 37°C, 5% CO_2_ before development. Development of the FluoroSpot plates was performed according to the manufacturer’s recommendations. After removing cells, detection antibodies diluted in PBS containing 0.1% BSA were added to the wells and the FluoroSpot plates were incubated for 2 hours at room temperature. Before read-out using the Mabtech IRIS™ automated FluoroSpot reader, the FluoroSpot plates were emptied and dried at room temperature for 24 h protected from light. All data were acquired with a Mabtech IRIS™ reader and analyzed using Mabtech Apex TM software. Unstimulated (DMSO) negative control, CEF positive control (T-cell epitopes derived from CMV, EBV and influenza, Mabtech, Sweden), and a commercial SARS-CoV-2 peptide pool (SARS-CoV-2 S N M O defined peptide pool (3622-1; Mabtech, Sweden) were included as assay controls. *Ex vivo* FluoroSpot results were considered positive when the test result was higher than DMSO negative control after subtracting non-stimulated control (dSFU).

Enriched FluoroSpot assays for convalescent donor testing were performed by Nexelis-IMXP (Belgium) as follows: PBMCs were retrieved from cryogenic storage and thawed in culture medium. The PBMCs were seeded at 4,000,000 cells/24-well in presence of the peptide pools (5 μg/ml per peptide) and incubated for 7 days at 37°C, 5% CO_2_. On days 1 and 4 of culture, the media were refreshed and supplemented with 5 ng/mL IL-7 or 5 ng/mL IL-7 and 4 ng/ml IL-2 (R&D Systems), respectively. After 7 days of culture, the PBMCs were harvested and rested for 16 h. The rested PBMCs were then counted using Trypan Blue Solution, 0.4% (VWR) and the Cellometer K2 Fluorescent Viability Cell Counter (Nexcelom), and seeded on the IFN-γ/Granzyme-B/TNF-α FluoroSpot plates (Mabtech) at 200,000 cells/well in RPMI 1640 with 10% Human Serum HI, 2 mM L-glutamine, 50 µg/mL gentamycin and 100 μM β-ME into the relevant FluoroSpot wells containing peptide (5 μg/mL), or peptide pool (5 μg/mL per peptide), in triplicates. The FluoroSpot plates were incubated overnight at 37°C, 5% CO_2_ before development. All data were acquired with a Mabtech IRIS™ reader and analyzed using Mabtech Apex TM software. DMSO, medium only, a commercial COVID peptide pool (SARS-CoV-2 S N M O defined peptide pool [3622-1] – Mabtech), and CEF were included as assay controls at a concentration of 1µg/ml. The positivity criterion was >1.5-fold the unstimulated control after subtracting the background (dSFU).

### Intracellular cytokine staining (ICS) assay

*Ex vivo* ICS assays for preclinical animal studies were performed as follows: splenocytes were removed from the ELISpot plates after 20 h of stimulation, transferred to a conventional 96-well flat bottom plate, and incubated for 4 h with BD GolgiStop^™^ according to the manufacturer’s recommendations. Flow-cytometry was performed using a BD Cytofix/Cytoperm Plus Kit with BD GolgiStop^™^ protein transport inhibitor (containing monensin; Cat. No. 554715), following the manufacturer’s instructions. Flow cytometry analysis and cytokine profile determination were performed on an Attune NxT Flow cytometer (Life Technologies). A total of 2 × 10^5^ cells were analyzed, gated for CD45^+^, CD3^+^, CD4^+^, or CD8^+^ T cells. Counts <25 were evaluated as 0. Spot counts ≥25 were background corrected by subtracting unstimulated (DMSO) control. PMA/Ionomycin (Invitrogen) was used as a positive control. As an assay control, Mann-Whitney test was used to compare negative control (unstimulated) and positive control (PMA/ionomycin) for each cytokine. When a statistical difference between controls was determined, the values corresponding to the other stimulation conditions were analyzed. The following stains were used for Hu-mice cohorts: MAb11 502932 (Biolegend), MP4-25D2 500836 (Biolegend), 4S.B3 502536 (Biolegend), HI30 304044 (Biolegend), SK7 344842 (Biolegend), JES6-5H4 503806 (Biolegend), VIT4 130-113-218 (Miltenyi), JES1-39D10 500904 (Biolegend), SK1 344744 (Biolegend), JES10-5A2 501914 (Biolegend), JES3-19F1 554707 (BD), and NA 564997 (BD). The following stains were used for BALB/c mice cohorts: 11B11 562915 (BD), MP6-XT22 506339 (Biolegend), XMG1.2 505840 (Biolegend), 30-F11 103151 (Biolegend), 145-2C11 100355 (Biolegend), JES6-5H4 503806 (Biolegend), GK1.5 100762 (Biolegend), JES1-39D10 500904 (Biolegend), 53-6.7 100762 (Biolegend), eBio13A 25-7133-82 (Thermo Scientific), JESS-16E3 505010 (Biolegend), and NA 564997 (BD).

*Ex vivo* ICS assays for convalescent donor testing were performed by Nexelis-IMXP (Belgium). Briefly, after thawing 200,000 PBMC cells/well, PBMCs were seeded in sterile round-bottom 96-well plates in RPMI total with 10% human serum HI, 2 mM L-glutamine, 50 μg/mL gentamycin, and 100 μM 2-ME in the presence of peptides (5 μg/mL) or peptide pool (5 μg/mL per peptide). After a 2-hour incubation, BD GolgiPlug^™^ (BD Biosciences) was added to the 96-well plates at a concentration of 1 μl/mL in culture medium. After a 10-h incubation, plates were centrifuged (800 × *g*, 3 min, 8°C) and incubated for 10 min at 37°C and Zombie NIR Viability dye (Biolegend) was added to each well. Plates were incubated at room temperature for 15 min, shielded from the light. After incubation, PBS/0.1% BSA was added per well and the plates were centrifuged (800 × *g*, 3 min, 8°C). Thereafter, cells were incubated in FcR blocking reagent at 4°C for 5 min, and then staining mixture (containing anti-CD3, Biolegend, anti-CD4, and anti-CD8 antibodies; BD Biosciences) was added to each well. After 30 min of incubation at 4°C, washing, and centrifugation (800 × *g*, 3 min, 8°C), cells were permeabilized and fixed according to the manufacturer’s recommendations (BD Biosciences). After fixation, cytokine staining mixture (containing anti-IFN-γ, anti-IL-2, anti-IL-4, anti-IL-10 and anti-TNF-α antibodies, Biolegend) was added to each well. Plates were incubated at 4°C for 30 min and then washed twice before acquisition. All flow cytometry data were acquired with LSRFortessa™ X-20 and analyzed using FlowJo V10 software. DMSO negative control was subtracted from each data point obtained using test peptides or pools.

### Antibody ELISA

ELISAs for mouse studies were performed for the quantitative measurement of total mouse IgG production in plasma samples using IgG (Total) Mouse Uncoated ELISA Kit (Invitrogen, #88-50400-22) for BALC/c cohorts and IgG (Total) Human Uncoated ELISA Kit (Invitrogen, #88-50550-22) for Hu-mice cohorts according to the manufacturer’s instructions. Analyses were performed using samples harvested at days 14, 21, and 28 (n=6 per group per time point). Absorbance were read on an Epoch Microplate Reader (Biotech) and analyzed using Gen5 software.

Euroimmune ELISA assays for convalescent donors were performed to determine S1-specific IgG levels via the Independent Medical Research Center, The Netherlands. The Anti-SARS-CoV-2 ELISA plates ere coated with recombinant S-1 structural protein from SARS-CoV-2 to which antibodies against SARS-CoV-2 bind. This antigen was selected for its relatively low homology to other coronaviruses, notably SARS-CoV. The immunoassay was performed according to the manufacturer’s instructions.

ELISAs were performed by Mikromikomed Kft (Budapest, Hungary) using a DiaPro COVID-19 IgM Enzyme Immunoassay for the determination of IgM antibodies to COVID-19 in human serum and plasma, DiaPro COVID-19 IgG Enzyme Immunoassay for the determination of IgG antibodies to COVID-19 in human serum and plasma, and DiaPro COVID-19 IgA Enzyme Immunoassay for the determination of IgA antibodies to COVID-19 in human serum and plasma according to the manufacturer’s instructions (Dia.Pro Diagnostic Bioprobes S.r.l., Italy). For the determination of N-specific antibodies, Roche Elecsys® Anti-SARS-CoV-2 Immunoassay for the qualitative detection of antibodies (including IgG) against SARS-CoV-2 was used with a COBAS e411 analyzer (disk system; Roche, Switzerland) according to the manufacturer’s instructions.

### Pseudoparticle Neutralization Assay (PNA)

Neutralizing antibodies in mice sera were assessed using a cell-based Pseudoparticle Neutralization Assay. Vero E6 cells expressing the ACE-2 receptor (Vero C1008 [ATCC No. CRL-1586, US]), were seeded at 20 000 cells/well to reach a cell confluence of 80%. Serum samples and controls (pool of human convalescent serum, internally produced) were diluted in duplicates in cell growth media at a starting dilution of 1/25 or 1/250 (for samples) or 1/100 (for controls) followed by a serial dilution (two-fold dilutions, five times). In parallel, SARS-CoV-2 pseudovirus (prepared by Nexelis, using Kerafast system, with S from Wuhan Strain (minus 19 C-terminal amino acids) was diluted to reach the desired concentration (according to pre-determined TU/mL). Pseudovirus was then added to diluted sera samples and pre-incubated for 1 hour at 37°C with CO_2_. The mixture was then added to the pre-seeded Vero E6 cell layers and plates were incubated for 18–24 hours at 37°C with 5% CO_2_. Following incubation and removal of media, ONE-Glo EX Luciferase Assay Substrate (Promega, Cat. E8110) was added to cells and incubated for 3 minutes at room temperature with shaking. Luminescence was measured using a SpectraMax i3x microplate reader and SoftMax Pro v6.5.1 (Molecular Devices). Luminescence results for each dilution were used to generate a titration curve using a 4-parameter logistic regression (4PL) using Microsoft Excel (for Microsoft Office 365). The titer was defined as the reciprocal dilution of the sample for which the luminescence is equal to a pre-determined cut-point of 50, corresponding to 50% neutralization. This cut-point was established using linear regression using 50% flanking points.

### Statistical analysis

Significance was compared between and among groups using *t*-tests, Mann-Whitney tests, or Permutation statistics using Montecarlo simulations as appropriate. p<0.05 was considered significant. Pearson’s test and or Spearman’s test was performed to assess correlations. The correlation was considered strong if R >0.71, moderate if R = 0.51–0.70 and weak if R = 0.31–0.50.(*54*)

## Results

### Tailoring PolyPEPI-SCoV-2 to individuals

During the design of PolyPEPI-SCoV-2, we used the HLA genotype data of subjects in the *in silico* human cohort to determine the most immunogenic peptides (i.e., HLA class I PEPI hotspots, 9-mers) of the four selected SARS-CoV-2 structural proteins aimed to induce CD8^+^ T cell responses. The sequences of the identified 9-mer PEPI hotspots were then extended to 30-mers based on the viral protein sequences to maximize the number of HLA class II binding PEPIs (15-mers) aimed to induce CD4^+^ T cell responses.

First, we performed epitope predictions for each subject in the *in silico* human cohorts for each of their HLA class I and class II genotypes (six HLA class I and eight HLA class II alleles) for the AA sequence of the conserved regions of 19 known SARS-CoV-2 viral proteins using 9-mer (HLA class I) and 15-mer (HLA class II) frames, respectively (Figure 1 and Methods). Then, we selected the epitopes that could bind to multiple (≥3) autologous HLA alleles (PEPIs) to account for the most abundantly presented epitopes on the cell surface. We identified several HLA-restricted epitopes (average, 166 epitopes only for S-1 protein) for each person. In contrast, PEPIs are represented at much lower level in all ethnicities (average, 14 multi-HLA-binding epitopes, Figure 1). Of note, we did not observe any difference in SARS-CoV-2-specific epitope generation capacity of individuals with different ethnicities based on their complete HLA-genotype, which does not seem to explain the higher infection and mortality rates observed in BAME.

**Figure 1.**
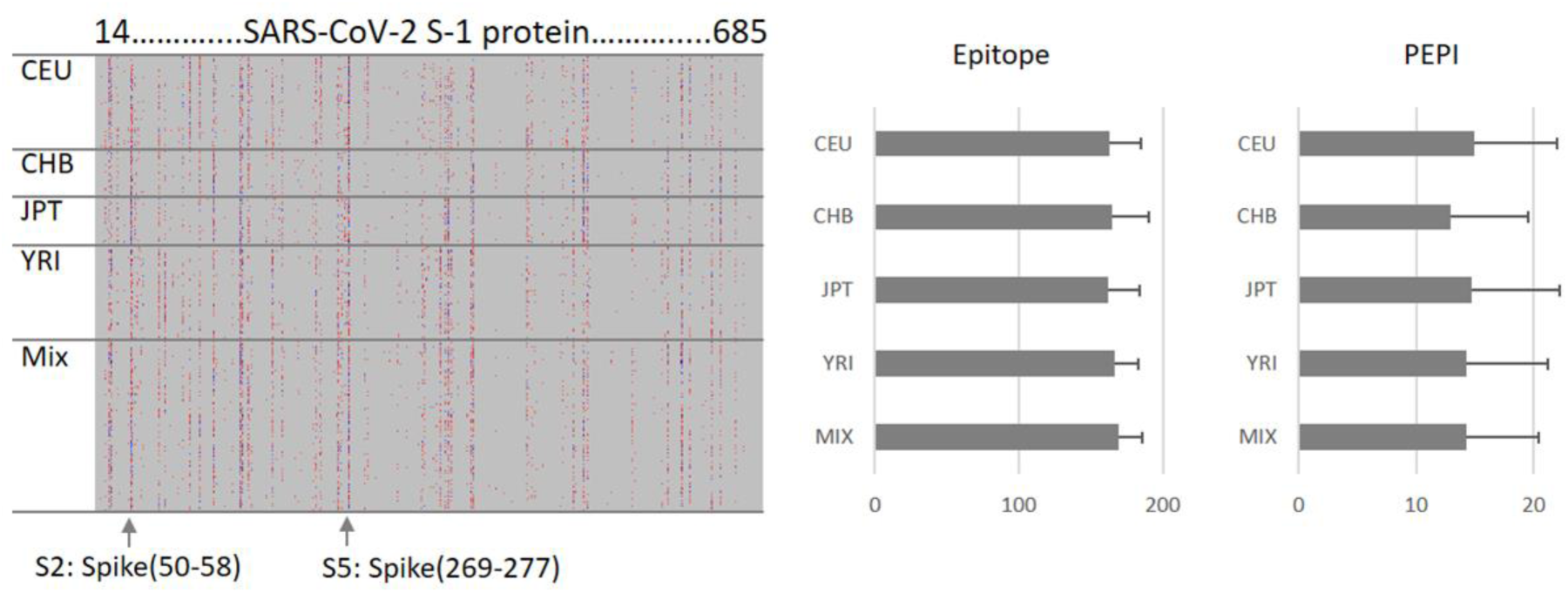
Hotspot analysis of SARS-CoV-2 Spike-1 protein in the ethnically diverse *in silico* human cohort. Analysis was performed by predicting ≥3 HLA allele binding personal epitopes (PEPIs) for each subject. Left panel: Each row along the vertical axis represents one subject in the model population, while the horizontal axis represents the SARS-CoV-2 S-1 protein sequence. Vertical bands represent the most frequent epitopes, i.e., the dominant immunogenic protein regions (hotspots) or PEPIs for most subjects. CEU, Central European; CHB, Chinese; JPT, Japanese; YRI, African; Mix, mixed ethnicity subjects. Colors represent the number of epitopes restricted to a person: red, 3; green/blue, 4; black, >5. A PEPI was defined as an epitope restricted to ≥3 alleles of a person. Right panel, average number of epitopes/PEPIs found for subjects of different ethnicities.

From the resulting PEPI list, we identified nine 30-mer polypeptide fragments that comprise overlapping, class I and class II PEPIs shared (frequent) among a high percentage of individuals in the model population, independent of ethnicity (Table 1). To maximize multi-antigenic immune responses at both the individual and population levels, we selected more than one peptide from the large S and N proteins and only one peptide from the shorter M and E proteins. From the four structural viral antigens of the SARS-CoV-2 virus, a total of nine 30-mer peptides were selected for the vaccine, also considering the chemical and manufacturability properties of the peptides: three peptides derived from S, four peptides from N, and one peptide derived from each M and E. No peptides were included from the receptor-binding domain of S-1 protein. Overall, each member of the model population had HLA class I PEPIs for at least two of the nine peptides, and 97% had at least three (Table 1). Each subject had multiple class II PEPIs for the vaccine peptides (Table 1)

We identified experimentally confirmed linear B cell epitopes derived from SARS-CoV, with 100% sequence identity to the relevant SARS-CoV-2 antigen, to account for the potential B cell production capacity of the long peptides.(*58*) Three overlapping epitopes located in N protein- and one epitope in S-protein-derived peptides of PolyPEPI-SCoV-2 vaccine were reactive with the sera of convalescent patients with severe acute respiratory syndrome (SARS). This suggests that the above antigenic sites on the S and N protein are important in eliciting a humoral immune response against SARS-CoV and likely against SARS-CoV-2 in humans.

As expected, PolyPEPI-SCoV-2 contains several (eight out of nine) peptides that are cross-reactive with SARS-CoV due to high sequence homology between SARS-CoV-2 and SARS-CoV. Sequence similarity is low between the PolyPEPI-SCoV-2 peptides and common (seasonal) coronavirus strains belonging to alpha coronavirus (229E and NL63), beta coronavirus (OC43, HKU1), and MERS. Therefore, cross-reactivity between the vaccine and prior coronavirus-infected individuals remains low and might involve only M1 peptide of the vaccine (See Methods and Supplementary Table 2).

### PolyPEPI-SCoV-2 vaccine induced broad T cell responses in two animal models

Preclinical immunogenicity testing of PolyPEPI-SCoV-2 vaccine was performed to measure the induced immune responses after one and two vaccine doses that were administered two weeks apart (days 0 and 14) in BALB/c and Hu-mouse models. After immunizations, no mice presented any clinical score (score 0, representing no deviation from normal), suggesting the absence of any side effects or immune aversion. In addition, necropsies performed by macroscopic observation at each timepoint did not reveal any visible organ alteration in spleen, liver, kidneys, stomach, and intestine. Repeated vaccine administration was also well tolerated, and no signs of immune toxicity or other systemic adverse events were detected. Together, these data strongly suggest that PolyPEPI-SCoV-2 was safe in mice.

Vaccine-induced IFN-γ producing T cells were measured after the first dose at day 14 and after the second dose at days 21 and 28. Vaccine-induced T cells were detected using the nine 30-mer vaccine peptides grouped in four pools according to their source protein: S, N, M, and E, to assess for the CD4+ and CD8+ T cell responses. CD8^+^ T cell responses were measured using the short 9-mer test peptides corresponding to the shared HLA class I PEPIs defined for each of the nine vaccine peptides that were also grouped into four pools according their source protein (S, N, M, and E peptides; Table 1 bold).

In BALB/c mice at day 14, PolyPEPI-SCoV-2 treatment did not induce more IFN-γ production than Vehicle (DMSO/Water emulsified with Montanide) treatment, the latter resulting in an unusually high response probably due to Montanide-mediated non-specific responses. Nevertheless, at days 21 and 28, the second dose of PolyPEPI-SCoV-2 increased IFN-γ production compared with Vehicle control group by 6-fold and 3.5-fold for splenocytes stimulated with the 30-mer and 9-mer peptides, respectively (Figure 2A).

**Figure 2.**
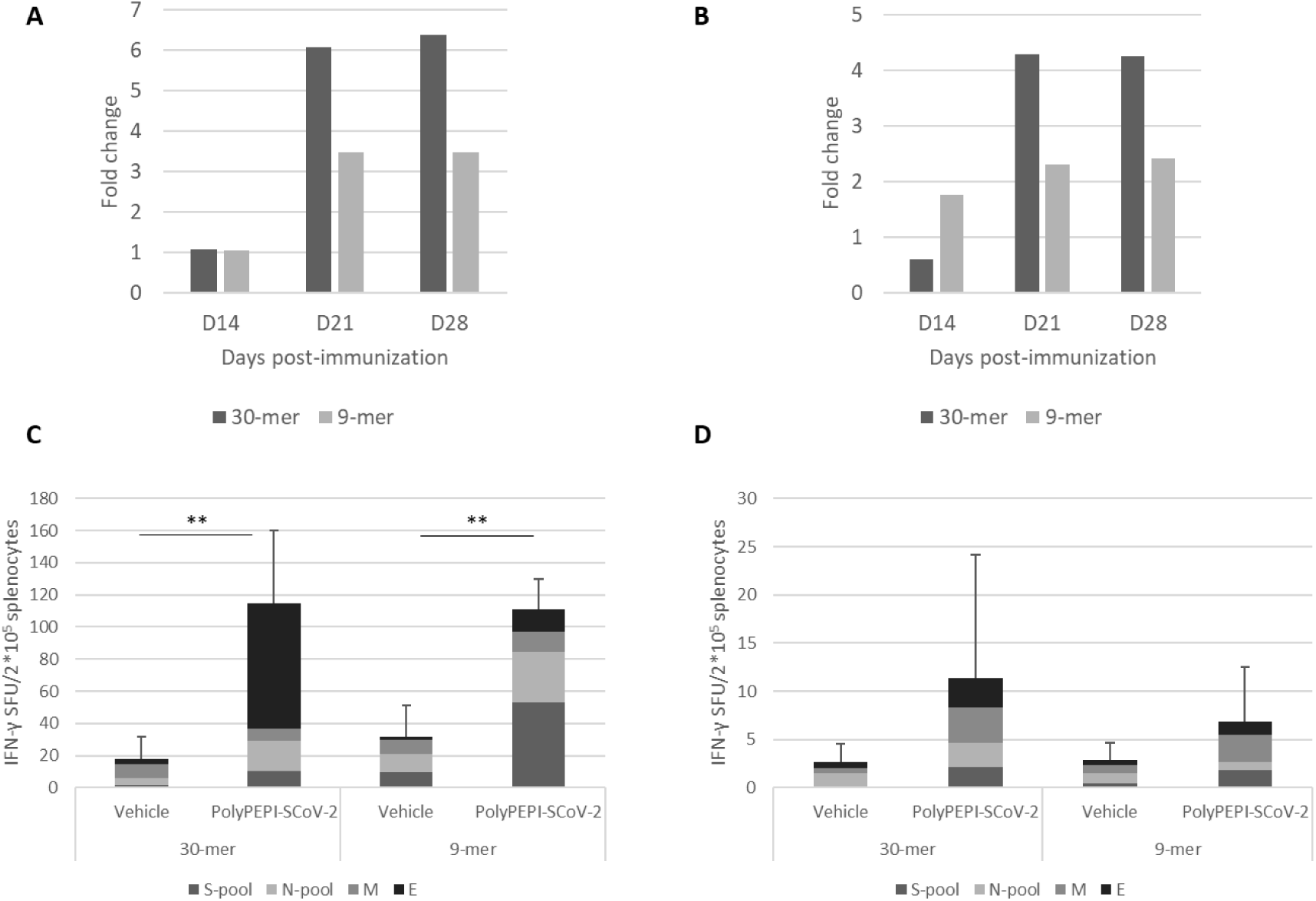
IFN-γ + T cell responses elicited by PolyPEPI-SCoV-2 vaccination in two animal models. Fold change in PolyPEPI-SCoV-2 vaccine-induced T cell responses in BALB/c mouse (**A**) and humanized (Hu-mouse) models (**B**) compared with the respective control cohorts receiving Vehicle only. Vaccine-induced T cell responses specific for SARS-CoV-2 protein-derived vaccine peptides after two doses detected at day 28 in BALB/c (**C**) and humanized (Hu-mice) (**D**). Mice received two doses of vaccine or Vehicle at days 0 and 14. Each cohort comprised six animals at each timepoint. *Ex vivo* ELISpot assays were performed by stimulation with 9-mer and 30-mer peptides. Test conditions: S-pool contains the three peptides derived from S protein; N-pool contains the four peptides derived from N protein; M and E are the peptides derived from M and E proteins, respectively, in both the 9-mer and 30-mer pools. Results were compared to Vehicle (DMSO/Water emulsified with Montanide) control group of the same time point. Spot forming unit (SFU) represents unstimulated background corrected values given for 2 × 10^5^ splenocytes. For fold change calculations, the mean values were obtained by pooling the background subtracted values of the four stimulation conditions for 30-mer (S-pool, N-pool, E1 and M1 peptides) and the four stimulation conditions for 9-mers (S-pool, N-pool, e1 and m1 peptides). t-tests were used to calculate significance. ^**^p<0.001.

In immunodeficient Hu-mice at day 14, PolyPEPI-SCoV-2 treatment increased IFN-γ production by two-fold with splenocytes stimulated with the 9-mer pool of peptides, but no increase was observed with 30-mer-stimulated splenocytes. At days 21 and 28, the second dose of PolyPEPI-SCoV-2 boosted IFN-γ production by two- and four-fold with splenocytes stimulated with the 9-mer and 30-mer pools of peptides, respectively (Figure 2B). Importantly, both 9-mer-detected CD8^+^ T cells and 30-mer-detected CD4^+^ and CD8^+^ T cell responses were directed against all four viral proteins targeted by the vaccine in both animal models (Figures 2C, D and Supplementary Figures 1A–F).

ICS assays was performed to investigate the polarization of the T cell responses elicited by the vaccination. Due to the low frequency of T cells, individual peptide-specific T cells were more difficult to visualize by ICS than by ELISpot, but a clear population of CD4^+^ and CD8^+^ T cells producing Th1-type cytokines of TNF-α and IL-2 were detectable compared with animals receiving only Vehicle in both BALB/c and Hu-mouse models (Figure 3). For IL-4 and IL-13 Th2-type cytokines, analysis did not reveal any specific response at any timepoint. Low levels of IL-5 and/or IL-10 cytokine-producing CD4^+^ T cells were detected for both models, but it was significantly different from Vehicle control only for BALB/c mice at day 28. Even for this cohort, the Th1/Th2 balance remained shifted toward Th1 for five out of six mice (one outlier), confirming an overall Th1-skewed T cell response elicited by the vaccine (Supplementary Figure 2).

**Figure 3.**
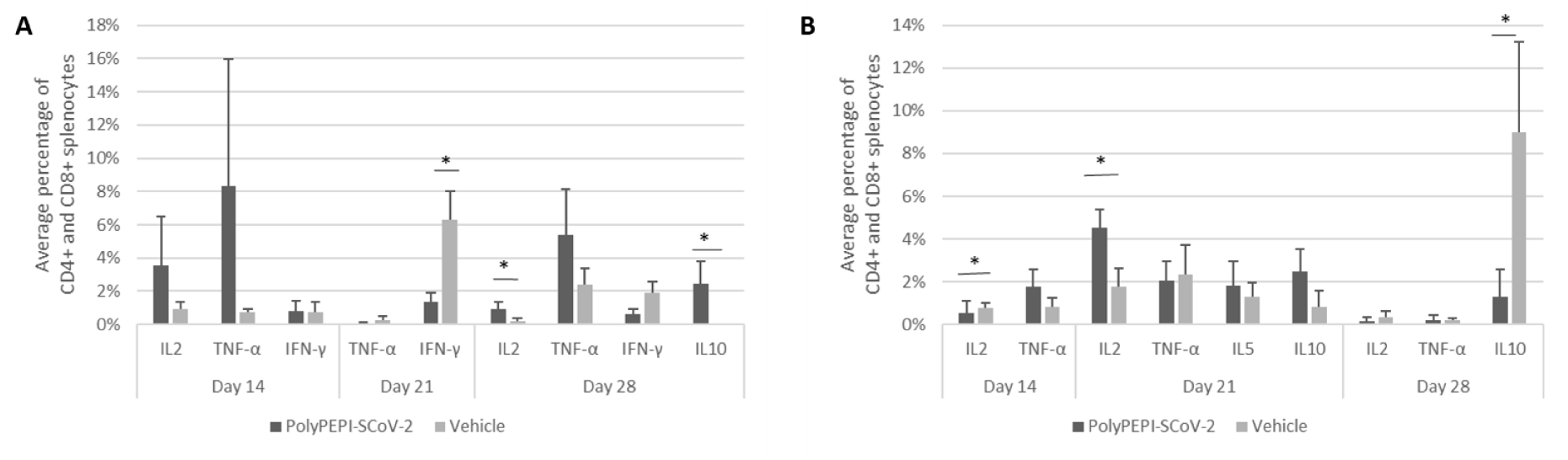
Th1 dominant immune response and no induction of significant Th2 cytokines with PolyPEPI-SCoV-2. Average CD4^+^ and CD8^+^ T cells producing IL2, TNF-α, IFN-γ, IL-5, or IL10 in immunized and Vehicle control groups for BALB/c (**A**) and Hu-mouse (**B**) models using ICS assays. Mean ± SEM are shown. 2 × 10^5^ cells were analyzed after gating for CD45^+^ cells, CD3^+^ T cells, CD4^+^, or CD8^+^ T cells. The mean percent was obtained by pooling the background subtracted values of the four stimulation conditions (30-mer S-pool, N-pool, E1 and M1 peptides) for each cytokine for CD4^+^ and CD8^+^ T cells.

PolyPEPI-SCoV-2 vaccination also induced humoral responses, as measured by total mouse IgG for BALB/c and human IgG for Hu-mouse models. In BALB/c mice, vaccination resulted in vaccine-induced IgG production after the first dose (day 14) compared with control animals receiving only Vehicle. IgG elevation was observed for both BALB/c and Hu-mouse models at later time points after the second dose (Figure 4). As expected, given that PolyPEPI-SCoV-2 peptides do not contain conformational B cell epitopes, vaccination did not result in neutralizing antibodies, as assessed from the sera of Hu-mice using PNA assays (data not shown).

**Figure 4.**
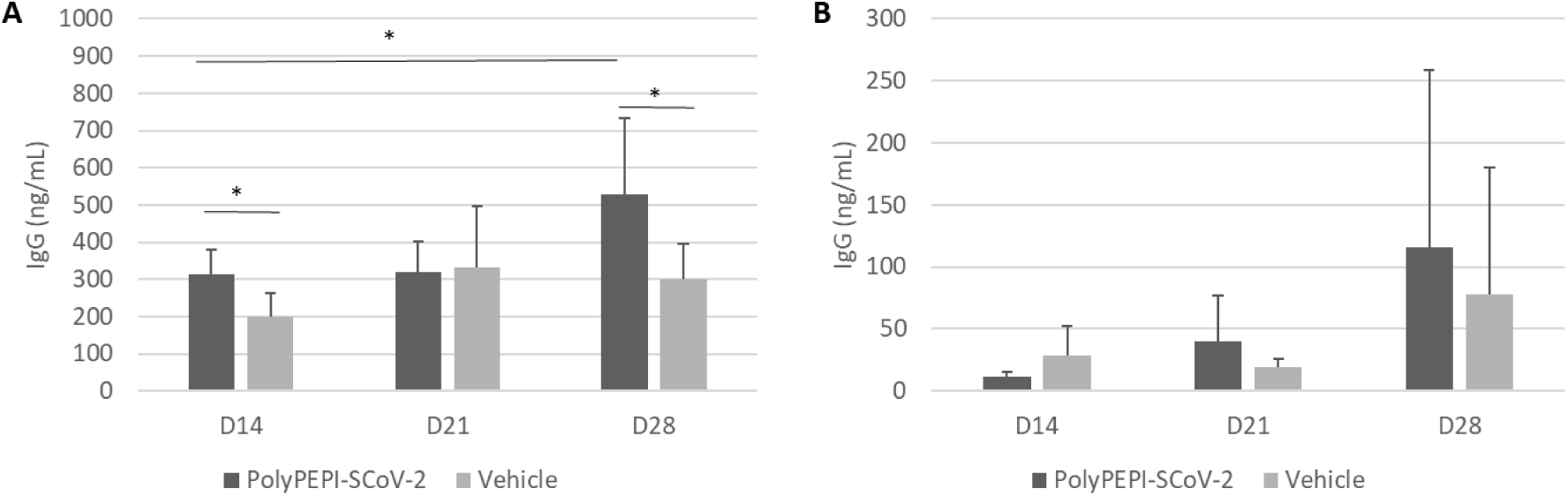
Vaccine-induced IgG production measured from the plasma of BALB/c mice (A) and Hu-mice (B). Mice received two doses of PolyPEPI-SCoV-2 vaccine or Vehicle at days 0 and 14. Each cohort comprised six animals. t-tests were used to calculate significance, p<0.05.

### PolyPEPI-SCoV-2-specific T cell responses detected in COVID-19 convalescent donors

Next, we aimed to demonstrate that the robust and broad PolyPEPI-SCoV-2-specific T cell responses detected in vaccinated animals are relevant in humans by investigating vaccine-specific T cells circulating in the blood of COVID-19 convalescent donors (baseline data are reported in Supplementary Table 1). First, the reactivity of vaccine peptides with convalescent immune components was investigated in 17 convalescent and four healthy donors using *ex vivo* FluoroSpot assays, which can detect rapidly activating, effector phase T cell responses. Vaccine-reactive CD4^+^ and CD8^+^ T cells were detected using the nine 30-mer vaccine peptides grouped in four pools according to their source protein: S, N, M, and E peptides. CD8^+^ T cell responses were measured using the 9-mer test peptides corresponding to the shared HLA class I PEPIs defined for each of the nine vaccine peptides that were also grouped into four pools according their source protein (S, N, M, and E peptides; Table 1 bold), as used in the animal experiments.

Significant amounts of vaccine-reactive, IFN-γ-expressing T cells were detected with both 30-mer (average dSFU: 48.1) and 9-mer peptides (average dSFU: 16.5) compared with healthy subjects (Figure 5A, B). Detailed analysis of the four protein-specific peptide pools revealed that three out of the 17 donors reacted to all four structural antigens with the 30-mer vaccine peptides; 82% of donors reacted to two antigens and 59% to three antigens. Notably, highly specific 9-mer detected CD8^+^ T cell responses could be also identified against at least one of four antigens in all 17 donors and against at least two antigens in 53% (Supplementary Table 3).

**Figure 5.**
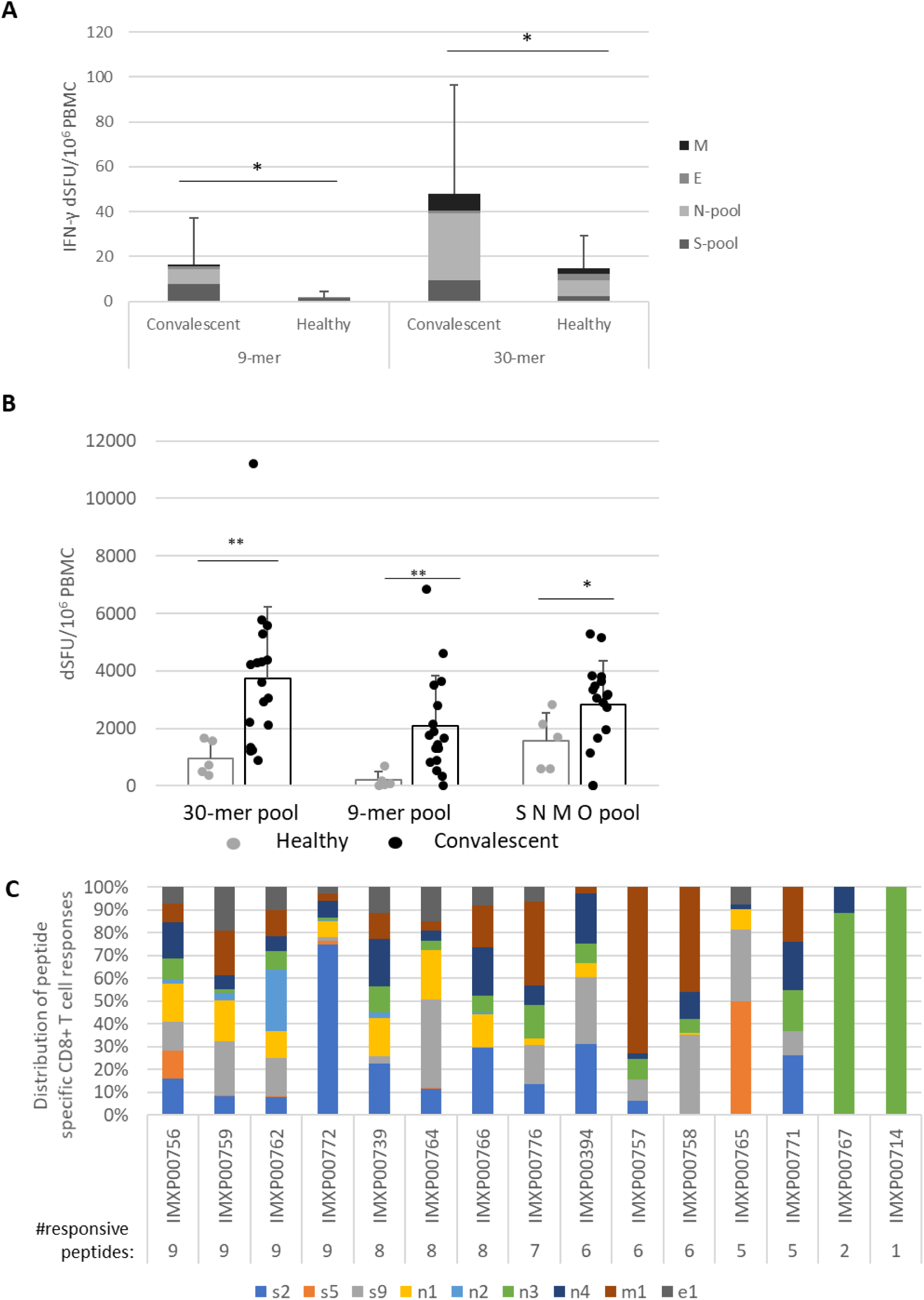
PolyPEPI-SCoV-2-specific T cell responses from COVID-19 convalescent donors. **A.** Highly specific vaccine-derived 9-mer peptide-reactive CD8^+^ T cell responses and 30-mer peptide-reactive CD4^+^ T cell responses detected by *ex vivo* FluoroSpot assay. **B**. Enriched FluoroSpot results with 30-mer peptide pools, 9-mer peptide pools, and commercial SNMO peptide pool-activated IFN-γ producing T cells. **C**. Enriched Fluorospot results with IFN-γ producing CD8^+^ T cells activated by individual 9-mer peptides corresponding to each of the 30-mer peptides with the same name (Table 1 bold). dSFU, delta spot forming units, calculated as non-stimulated background corrected spot counts per 10^6^ PBMC. Significance was calculated using Permutation statistics with Montecarlo simulations; ^*^p<0.05, ^**^p<0.00005.

As determined by ICS assays, stimulation with 9-mer peptides resulted in an average T cell make up of 83% CD8^+^ T cells, and 17% CD4^+^ T cells (Supplementary Figure 3A). The 30-mer test peptides reacted with both CD4^+^ and CD8^+^ T cells in average ratio of 50:50 (Supplementary Figure 3A). Functionality testing of the T cells revealed that CD8^+^ T cells primarily produced IFN-γ, TNF-α, and IL-2 (with small amounts of IL-4 and IL-10), while CD4^+^ T cells were positive for mainly IL-2 and IFN-γ. Recall responses demonstrated clear Th1 cytokine characteristics; Th2 responses were not present in the recall response with 30-mer vaccine peptides (Supplementary Figure 3B).

Next, we determined whether the *ex vivo* detected T cells could also expand *in vitro* in the presence of vaccine peptides. Using enriched FluoroSpot, significant numbers of vaccine-reactive, IFN-γ-expressing T cells were detected with both 30-mer (average dSFU=3,746) and 9-mer (average dSFU=2,088) peptide pools compared with healthy subjects (Figure 5A). The intensity of the PolyPEPI-SCoV-2-derived T cell responses (30-mer pool) were also evaluated relative to the responses detected with a commercial, large SARS-CoV-2 peptide pool (SMNO) containing 47 long peptides derived from both structural (S, M, N) and non-structural (open reading frame ORF-3a and 7a) proteins. COVID-19 donors had a greater response to the vaccine pool, whereas healthy donors had a greater response to the commercial peptide pool, confirming improved specificity of PolyPEPI-SCoV-2 vaccine to SARS-CoV-2 (Figure 5B).

To confirm and further delineate the multi-specificity of the PolyPEPI-SCoV-2-specific T cell responses of COVID-19 recovered individuals, we defined the distinctive peptides targeted by their T cells. We first deconvoluted the peptide pools and tested the CD8^+^ T cell responses specific to each of the 9-mer HLA class I PEPIs corresponding to each vaccine peptide using *in vitro* expansion (Supplementary Figure 4). Analysis revealed that each 9-mer peptide was recognized by several subjects; the highest recognition rate in COVID-19 convalescent donors was observed for N4 and N3 (93%), S9 (87%), S2, N1, M1 (80%), E1 (60%), and finally S5 and N2 (40%) (Figure 5C). Detailed analysis of the nine peptide-specific CD8^+^ T cell responses revealed that 100% of COVID-19-recovered subjects had PolyPEPI-SCoV-2-specific T cells reactivated with at least one peptide, 93% with more than two, 87% with more than five, and 27% had T cell pools specific to all nine vaccine peptides. At the protein level, 87% of subjects had T cells against multiple (three) proteins and eight out of the 15 measured donors (53%) reacted to all four targeted viral proteins (Figure 5C). These data confirm that PolyPEPI-SCoV-2-peptides are dominant for an individual and shared between COVID-19 subjects, and that the peptide-specific T cell frequencies obtained in the convalescent population were in good alignment with the predicted frequencies based on shared PEPIs for the *in silico* cohort (Table 1). Convalescents’ T cells recognizing PolyPEPI-SCoV-2 specific 9-mer peptides were fully functional, expressing IFN-γ and/or TNF-α and/or Granzyme-B (Supplementary Figure 5). For our cohort of convalescent subjects, the breadth and magnitude of vaccine-specific T cell responses were independent of time from symptom onset, suggesting that these T cells are persistent (for at least 5 months) (Supplementary Figure 6).

To demonstrate that PEPIs (multi-HLA binding epitopes) can generate CD8^+^ T cell responses at the individual level, we first determined the complete class I HLA genotype for each subject and then predicted the peptides that could bind to at least three HLA alleles of a person from the list of nine shared HLA class I 9-mer peptides determined for the *in silico* cohort and used in the FluoroSpot assay. For each subject, between two and seven peptides out of nine were PEPIs. Among the predicted peptides (PEPIs), 84% were confirmed by IFN-γ FluoroSpot to generate highly specific T cell responses in the given subject (Table 2).

**Table 2.**
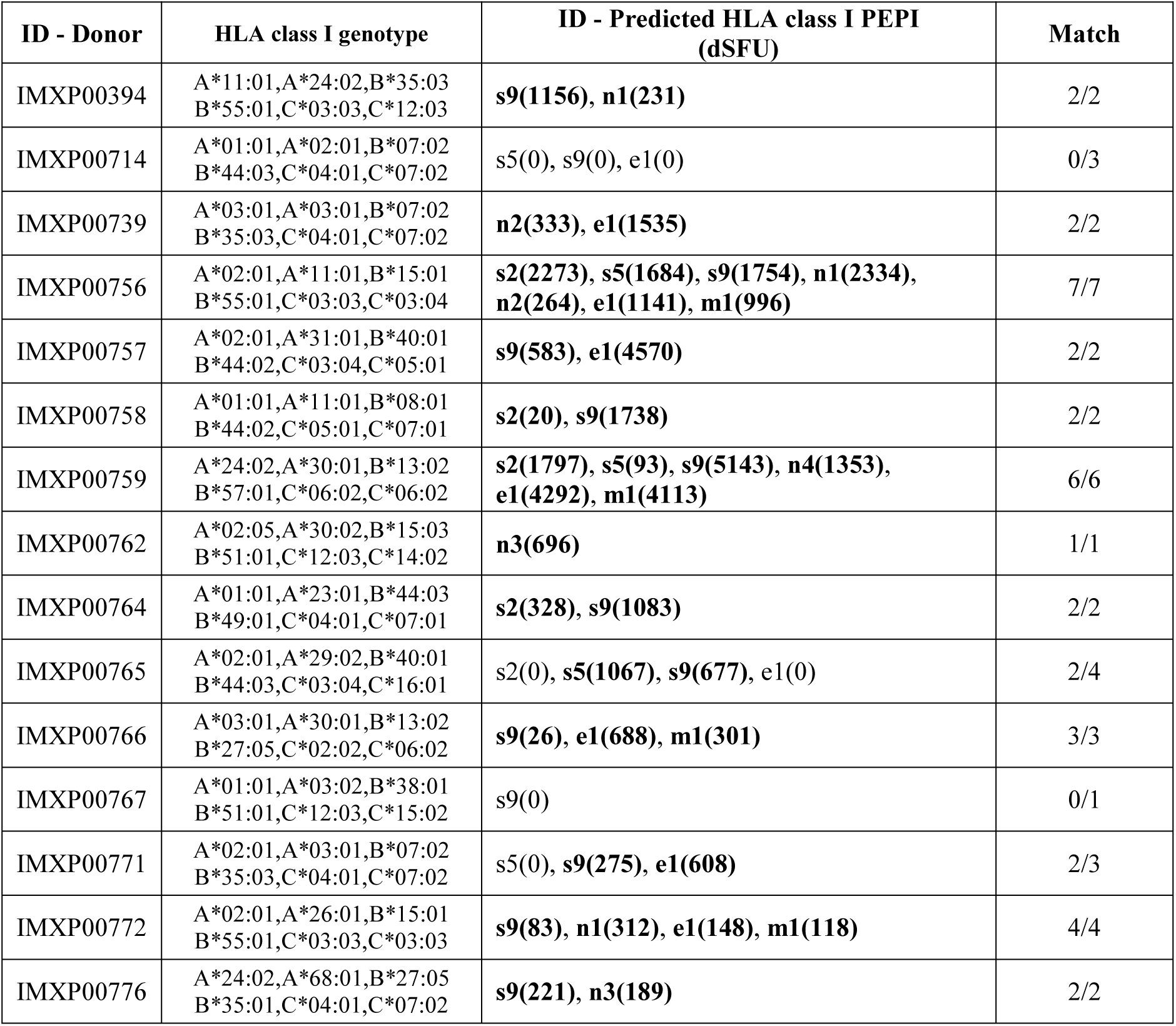
Agreement between PEPI prediction and immune responses measured by enriched ELISpot assay for COVID-19 convalescents. Personal epitopes (PEPIs) were predicted as ≥3 autologous HLA class I allele binding 9-mer epitopes and compared with the IFN-γ-producing CD8^+^ T cell response measured with identical 9-mer stimulations. True positive values are highlighted in bold letters.

| ID - Donor | HLA class I genotype | ID - Predicted HLA class I PEPI (dSFU) | Match |
| --- | --- | --- | --- |
| IMXP00394 | A*11:01,A*24:02,B*35:03<br>B*55:01,C*03:03,C*12:03 | **s9(1156), n1(231)** | 2/2 |
| IMXP00714 | A*01:01,A*02:01,B*07:02<br>B*44:03,C*04:01,C*07:02 | s5(0), s9(0), e1(0) | 0/3 |
| IMXP00739 | A*03:01,A*03:01,B*07:02<br>B*35:03,C*04:01,C*07:02 | **n2(333), e1(1535)** | 2/2 |
| IMXP00756 | A*02:01,A*11:01,B*15:01<br>B*55:01,C*03:03,C*03:04 | **s2(2273), s5(1684), s9(1754), n1(2334),**<br>**n2(264), e1(1141), m1(996)** | 7/7 |
| IMXP00757 | A*02:01,A*31:01,B*40:01<br>B*44:02,C*03:04,C*05:01 | **s9(583), e1(4570)** | 2/2 |
| IMXP00758 | A*01:01,A*11:01,B*08:01<br>B*44:02,C*05:01,C*07:01 | **s2(20), s9(1738)** | 2/2 |
| IMXP00759 | A*24:02,A*30:01,B*13:02<br>B*57:01,C*06:02,C*06:02 | **s2(1797), s5(93), s9(5143), n4(1353),**<br>**e1(4292), m1(4113)** | 6/6 |
| IMXP00762 | A*02:05,A*30:02,B*15:03<br>B*51:01,C*12:03,C*14:02 | **n3(696)** | 1/1 |
| IMXP00764 | A*01:01,A*23:01,B*44:03<br>B*49:01,C*04:01,C*07:01 | **s2(328), s9(1083)** | 2/2 |
| IMXP00765 | A*02:01,A*29:02,B*40:01<br>B*44:03,C*03:04,C*16:01 | s2(0), **s5(1067), s9(677)**, e1(0) | 2/4 |
| IMXP00766 | A*03:01,A*30:01,B*13:02<br>B*27:05,C*02:02,C*06:02 | **s9(26), e1(688), m1(301)** | 3/3 |
| IMXP00767 | A*01:01,A*03:02,B*38:01<br>B*51:01,C*12:03,C*15:02 | s9(0) | 0/1 |
| IMXP00771 | A*02:01,A*03:01,B*07:02<br>B*35:03,C*04:01,C*07:02 | s5(0), **s9(275), e1(608)** | 2/3 |
| IMXP00772 | A*02:01,A*26:01,B*15:01<br>B*55:01,C*03:03,C*03:03 | **s9(83), n1(312), e1(148), m1(118)** | 4/4 |
| IMXP00776 | A*24:02,A*68:01,B*27:05<br>B*35:01,C*04:01,C*07:02 | **s9(221), n3(189)** | 2/2 |

### Correlation between PolyPEPI-SCoV-2-reactive T cells and SARS-CoV-2-specific antibody responses

T cell-dependent B cell activation is required for antibody production. For each subject, different levels of antibody responses were detected against both S and N antigens of SARS-CoV-2 determined using different commercial kits (Supplementary Table 1). All subjects tested positive with Euroimmune ELISA against viral S-1 (IgG S1) and a Roche kit to measure N-related antibodies (IgG-N). All subjects tested positive for DiaPro IgG and IgM (except 2 donors), 7/17 for DiaPro IgA detecting mixed S-1 and N protein-specific antibody responses (Supplementary Table 1).

We next evaluated the correlation between PolyPEPI-SCoV-2-specific CD4^+^ T cell reactivities and antibody responses (Figure 6). The total amount of PolyPEPI-SCoV-2-reactive CD4^+^ T cells correlated with IgG-S1 (R=0.59, p=0.02, Figure 6A). Next, the subset of CD4^+^ T cells reactive to specific S-1 protein-derived peptides of the PolyPEPI-SCoV-2 vaccine (S2 and S5) were analyzed and the correlation was similar (R=0.585, p=0.02, Figure 6B). T cell responses detected with N protein derived PolyPEPI-SCoV-2 peptides (N1, N2, N3 and N4) presented a weak but not significant correlation with IgG-N (Figure 6C). These data suggest a link between PolyPEPI-SCoV-2-specific CD4^+^ T cell responses and subsequent IgG production for COVID-19 convalescent donors. Interestingly, IgA production correlated with PolyPEPI-SCoV-2-specific memory CD4^+^ T cell responses (R=0.63, p=0.006, Figure 6D, although Spearman test did not confirm the correlation). T cell responses reactive to the SMNO peptide pool exhibited no correlation with any of the antibody subsets. This suggests that not all CD4^+^ T cells contributed to B cell responses and consequently to IgG production (data not shown).

**Figure 6.**
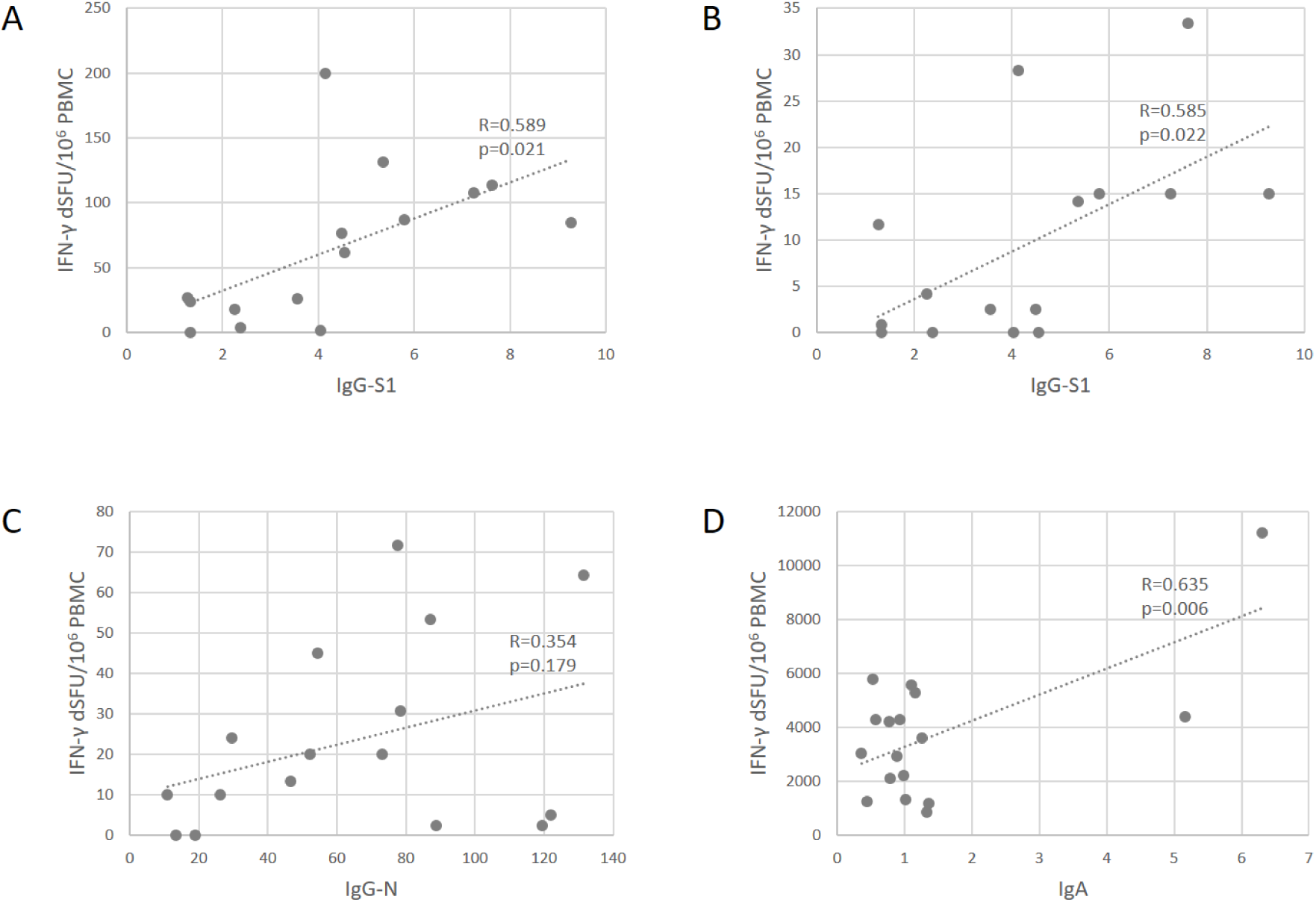
Correlation between SARS-CoV-2-specific antibody levels and PolyPEPI-SCoV-2-specific IFN-γ-producing CD4^+^ T cells in COVID-19 convalescent individuals. **A)** T cell responses reactive to 30-mer pool of PolyPEPI-SCoV-2 peptides were plotted against the IgG-S1 (Euroimmune). **B)** Average T cell responses reactive to S-1 protein-derived 30-mer peptides (S2 and S5) was plotted against IgG-S1 (Euroimmune). **C)** T cell responses reactive to 30-mer N peptide pool comprising N1, N2, N3 and N4 was plotted against total IgG-N measured with Roche Elecsys® assay. **D)** T cell responses reactive to 30-mer pool of PolyPEPI-SCoV-2 peptides were plotted against the IgA antibody amounts measured by DiaPro IgA ELISA assay. R-Pearson correlation coefficient.

### Predicted immunogenicity in different ethnicities

We performed *in silico* testing of our PolyPEPI-SCoV-2 vaccine in a large cohort of 16,000 HLA-genotyped subjects distributed among 16 different ethnic groups obtained from a US bone marrow donor database.(*53*) For each subject in this large cohort, we predicted the PolyPEPI-SCoV-2-specific PEPIs based on their complete HLA class I and class II genotype. Most subjects have a broad repertoire of PEPIs that will likely be transformed to virus-specific memory CD8^+^ T cell clones: 98% of subjects were predicted to have PEPIs against at least two vaccine peptides, and 95%, 86%, and 70% against three, four, and five peptides, respectively (Figure 7A).

**Figure 7.**
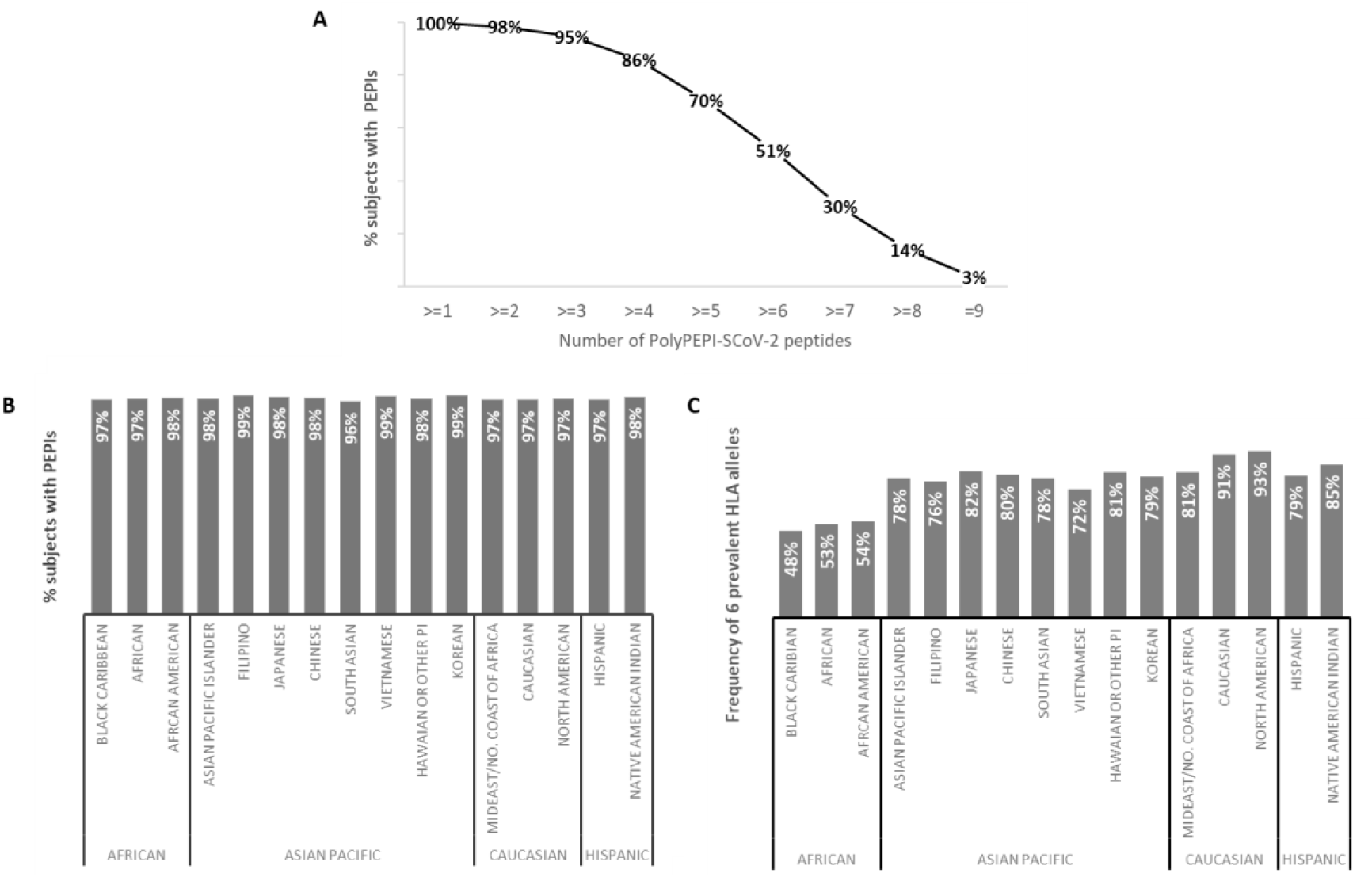
Predicted global coverage in a large population with different ethnicities. **A)** Proportion of subjects having HLA class I PEPIs against at least one of the nine PolyPEPI-SCoV-2 vaccine peptides **B)** Proportion of subjects having both HLA class I and class II PEPIs against at least two peptides in the PolyPEPI-SCoV-2 vaccine. **C)** Theoretical global coverage estimated based on the frequency of six prevalent HLA alleles (A^*^02:01, A^*^01:01,A^*^03:01, A^*^11:01, A^*^24:02, and B^*^07:02), as proposed by Ferretti et al.(*30*).

*In silico* testing revealed that 96-99% of subjects in each ethnic group will likely mount robust cellular responses, with both CD8^+^ and CD4^+^ T cell responses against at least two peptides in the vaccine (Figure 7B). This predicted high response rate is also true for the ethnicities reported to have worse clinical outcomes from COVID-19 (BAME).(*28*) Based on these data, we expect that the vaccine will provide global coverage, independent of ethnicity and geographic location.

We also used this large US cohort (and comprising ethnic groups) to assess theoretical global coverage as proposed by others, i.e., filtering the sub-populations having at least one of the six prevalent HLA class I alleles considered to cover 95% of the global population.(*30, 59, 60*) Using this approach, we observed significant heterogeneity at the ethnicity level. While we confirmed that the selected six HLA alleles are prevalent in the Caucasian and North American cohorts (91-93%), the frequency of these alleles was lower in all other ethnic groups, especially in African populations (48–54%) (Figure 7C). We concluded that the proposed prevalent HLA allele set may cover the HLA frequency in an ethnically weighted global population, but epitope selection for vaccination purposes based only on these alleles would discriminate some etnnicities. Therefore, we propose using a representative model population that is sensitive to the heterogeneities in the human race and that allows selecting PEPIs shared among individuals across ethnicities.

## Discussion

We demonstrated that PolyPEPI-SCoV-2, a polypeptide vaccine comprising nine synthetic long (30-mer) peptides derived from the four structural proteins of the SARS-CoV-2 (S, N, M, E) is safe and highly immunogenic in BALB/c mice and humanized CD34^+^ mice when administered with Montanide ISA 51 VG adjuvant. In addition, the vaccine’s immunogenic potential was confirmed in COVID-19 convalescent donors by successfully reactivating PolyPEPI-SCoV-2-specific T cells, which broadly overlap with the T cell immunity generated by SARS-CoV-2 infection.

Our vaccine design concept, targeting multi-antigenic immune responses at both the individual and population level, represents a novel target identification process that has already been used successfully in cancer vaccine development to achieve unprecedented immune response rates that correlate with initial efficacy in the clinical setting.(*35*) For COVID-19, we focused on selecting fragments of the SARS-CoV-2 proteins that contain overlapping HLA class I and II T cell epitopes shared between ethnically diverse HLA-genotyped individuals and that also generate diverse and broad immune responses against the whole virus structure. Therefore, we selected long 30-mer fragments to favor generation of multiantigenic effector responses (B cells and cytotoxic T cells) and helper T cell responses.

PolyPEPI-SCoV-2 vaccine elicits the desired multi-antigenic IFN-γ producing T cell responses of both vaccine-specific CD8^+^ and CD4^+^ T cells in vaccinated BALB/c and humanized CD34^+^ mice against all four SARS-CoV-2 proteins. The recall responses in COVID-19 convalescents comprised both rapidly activating effector-type (*ex vivo* detected) and expanded (*in vitro* detected) memory-type CD8^+^ and CD4^+^ T cell responses against all nine peptides, with PolyPEPI-SCoV-2-specific T cells detected in 100% of donors. At the individual level, the PolyPEPI-SCoV-2-specific T cell repertoire used for recovery from asymptomatic/mild COVID-19 was extremely diverse: each donor had an average of seven different peptide-specific T cell pools, with multiple targets against SARS-CoV-2 proteins; 87% of donors had multiple targets against at least three SARS-CoV-2 proteins and 53% against all four, 1–5 months after their disease. Although 87% of subjects had CD8^+^ T cells against S protein, which is similar with the immune response rates reported for frontline COVID-19 vaccine candidates in phase I/II clinical trials,(*61*) we found that S-specific (memory) T cells represented only 36% of the convalescents’ total T cell repertoire detected with vaccine peptides; the remaining 64% was distributed almost equally among N, M, and E proteins. These data support the increasing concern that S-protein based candidate vaccines are not harnessing the full potential of human anti-SARS-CoV-2 immunity, especially since diversity of T cell responses was associated with mild/asymptomatic COVID-19.*(4*)

The interaction between T and B cells is a well-known mechanism toward both antibody-producing plasma cell production and generation of memory B cells.(*62*) During the analysis of convalescents’ antibody subsets, we found correlations between antigen-specific IgG levels and corresponding peptide-specific CD4^+^ T cell responses. This correlation might represent the link between CD4^+^ T cells and antibody production, a concept also supported by total IgG production in animal models. Binding IgG antibodies can act in cooperation with the vaccine-induced CD8^+^ killer T cells upon later SARS-CoV-2 exposure of vaccinees. This interplay might result in effective CD8^+^ T cell mediated direct killing of infected cells and IgG-mediated killing of virus-infected cells and viral particles, inhibiting Th2-dependent immunopathologic processes. In this way, it is expected that both intracellular and extracellular virus reservoirs are attacked to help rapid viral clearance, even in the absence of neutralizing antibodies.(*62, 63*)

We demonstrated that individuals’ anti-SARS-CoV-2 T cell responses reactive to the PolyPEPI-SCoV-2 peptide set are HLA genotype-dependent. Specifically, multiple autologous HLA binding epitopes (PEPIs) determine antigen-specific CD8^+^ T cell responses with 84% accuracy. Although PEPIs generally underestimated the subject’s overall T cell repertoire, they are precise target identification “tools” and predictors of PEPI-specific immune responses, overcoming the high false-positive rates generally observed in the field using only the epitope-binding affinity as the T cell response predictor.(*37, 64*) Therefore, by means of validated PEPI prediction of T cell responses based on the complete HLA genotype of (only) Caucasian individuals and careful interpretation of PolyPEPI-SCoV-2-induced immune responses in animals as models for immune responses in humans, our findings could be extrapolated to large cohorts of 16,000 HLA-genotyped individuals and 16 human ethnicities.

Based on this, PolyPEPI-SCoV-2 will likely generate meaningful, multi-antigenic immune responses in ~98% of the global population, independent of ethnicity. In comparison, a T cell epitope-based vaccine design approach based on globally frequent HLA alleles would miss generation of immune responses for ~50% of Black Caribbean, African, African-American, and Vietnamese ethnicities. Inducing meaningful immune responses by vaccination (broad and polyfunctional memory responses mimicking the heterogeneity of the immunity induced by SARS-CoV-2) uniformly in high percentages of vaccinated subjects is essential to achieve the desired “herd immunity”. It is considered that about 25–50% of the population would have to be immune to the virus to achieve suppression of community transmission.(*3*) However, first-generation COVID-19 vaccines are being tested to show disease risk reduction of at least 50% but they are not expected to reduce virus transmission to a comparable degree. This fact combined with the likelihood that these vaccines will also not provide long-term immunity suggest that second-generation tools are needed to fight the pandemic.(*3*)

We believe that focusing on several targets in each subject would better recapitulate the natural T cell immunity induced by the virus, leading to efficient, long-term memory responses. For this purpose, synthetic polypeptide-based platform technology is considered a safe and immunogenic vaccination strategy with several key advantages. The same class of peptide vaccines with Montanide adjuvant as well as our PolyPEPI1018 vaccine with the same formulation were safe and well-tolerated in > 6,000 patients and 231 healthy volunteers to date.(*65–72*) In addition, efficient Th1-skewed T cell responses and non-conformational (linear) B cell epitopes mitigate the risk of theoretical antibody-mediated disease enhancement and Th2-based immunopathologic processes. Peptide-based vaccines have had only limited success to date, but this can be attributed to a lack of knowledge regarding which peptides to use. Such uncertainty is reduced by an understanding of how an individual’s genetic background can respond to specific peptides, as demonstrated above.

In conclusion, our peptide-based, multiiantigen-targeting vaccine, by demonstrating safety and exceptionally broad preclinical immunogenicity, has the potential to provide an effective second-generation solution against SARS-CoV-2, following careful clinical testing.

## Acknowledgement

We are grateful to all COVID-19 convalescent donors for the biospecimens; to Gabor Illes (Treos Bio Group) and to Florent Arbogast (TransCure Bioservices) who supported and facilitated the timely execution of the experiments.

## Author contributions

ES designed and coordinated the preclinical experiments, participated in data evaluations; ZC designed the PolyPEPI-SCoV-2 vaccine, participated in preclinical data evaluations; LM and JT performed the *in silico* analyses and prepared the Figures and Tables of the manuscript. SP, JS and AM performed the in vitro experiments using COVID-19 donors’ specimen and participated in the analysis of these data; OL had leading role in the manufacturing and quality control of the PolyPEPI-SCoV-2 peptides and development of vaccine formulation; LM, IM performed the statistical analysis; KP, PP participated in data mining and literature search; ERT participated in the design of the experiments and interpretation of data as well in the preparation of the manuscript. All authors reviewed the manuscript.

---

**Supplementary Table 1.**
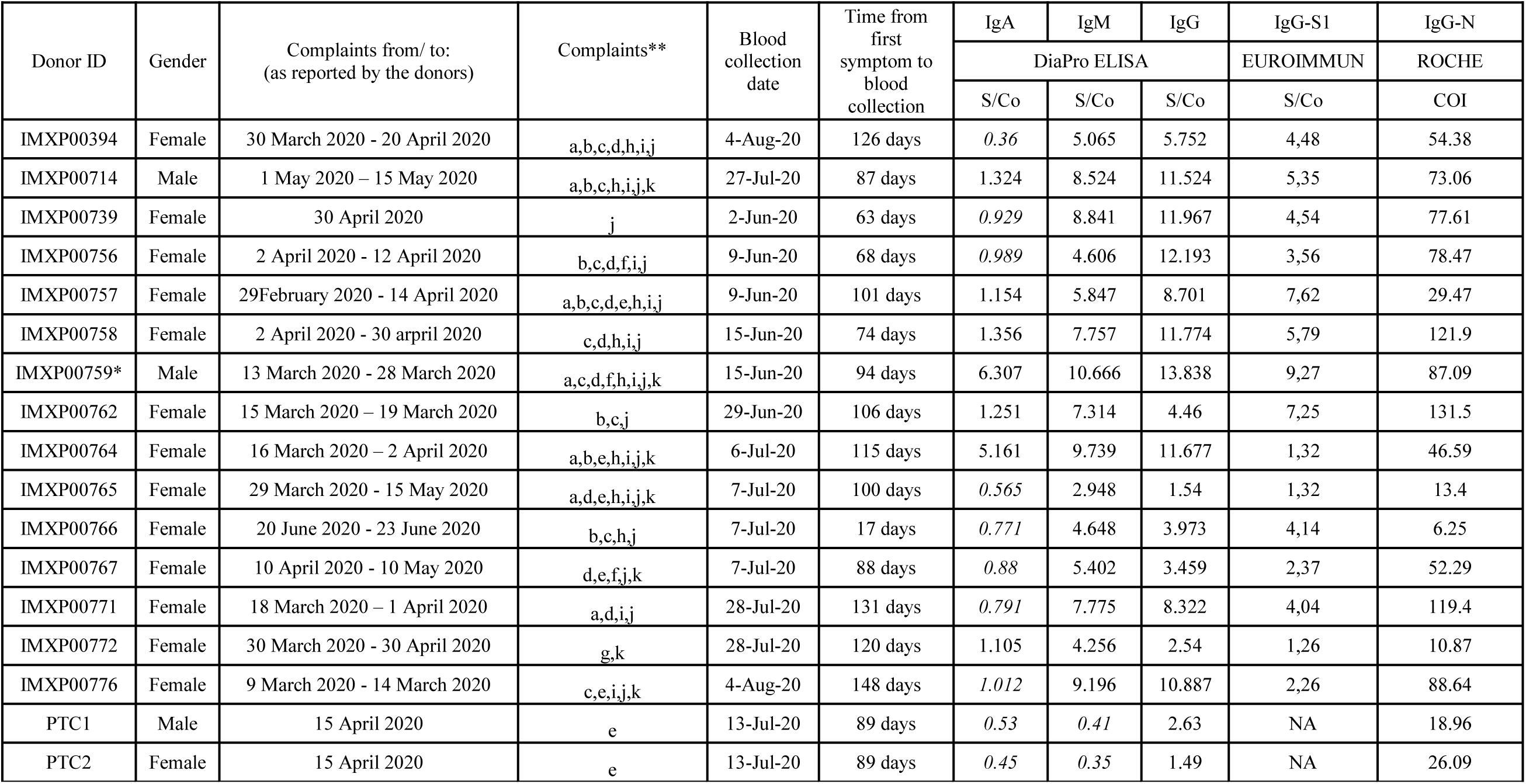
Donor baseline and demographic information. All donors were caucasoid, with mild/asymptomatic disease and no hospitalization (except one, marked with ^*^). S/Co, sample/control ratio; values were determined according to the manufacturer’s instructions, and test results are interpreted as negative in S/Co <0.9, not conclusive if S/CO = 0.9–1.1, and positive if S/Co >1.1. COI, cut-off index; values were determined according to the manufacturer’s instructions, and test results are interpreted as negative in COI <0.9, inconclusive with COI 0.9–1.1, and positive if COI >1.1. NA, data not available. Italic, negative or inconclusive values. ** Complaints: a, cough; b, sore throat; c, fever; d, short of breath; e, stomach/intestinal complaints; f, chest pain; g, sore eyes; h, odor or taste loss; i, headache; j, fatigue; k, other complaints (pulmonary embolism and cardiac arrest for IMXP00759; leg pain, arm pain, muscle pain, pain in the eyes).

**Supplementary Table 2.**
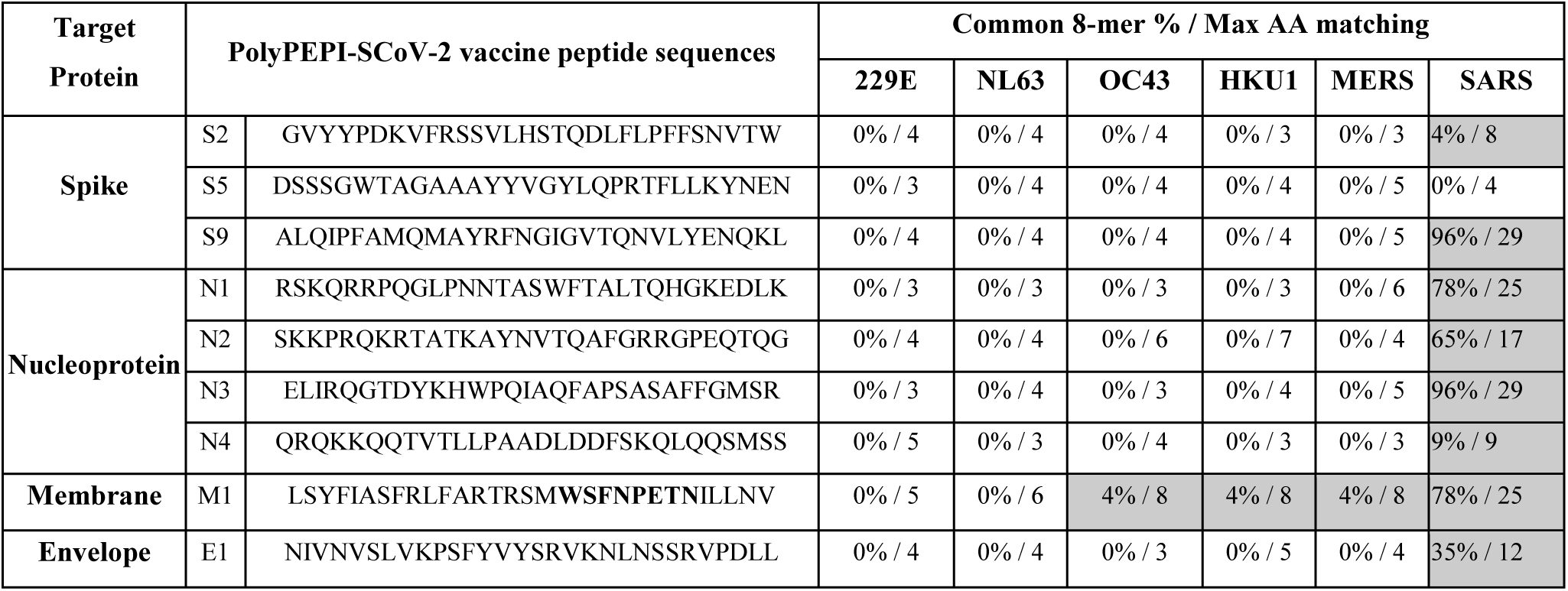
Sequence alignment results between PolyPEPI-SCoV-2 and other coronavirus strains. Sequence comparison was made with 8-mer long peptide matching between the aligned protein sequence pairs, defined as the minimum length requirement for a CD8^+^ T cell epitope. Max AA matching: the longest identical amino acid sequence length. Highlighted grey values represent identical sequences of at least eight amino acids.

**Supplementary Table S3.**
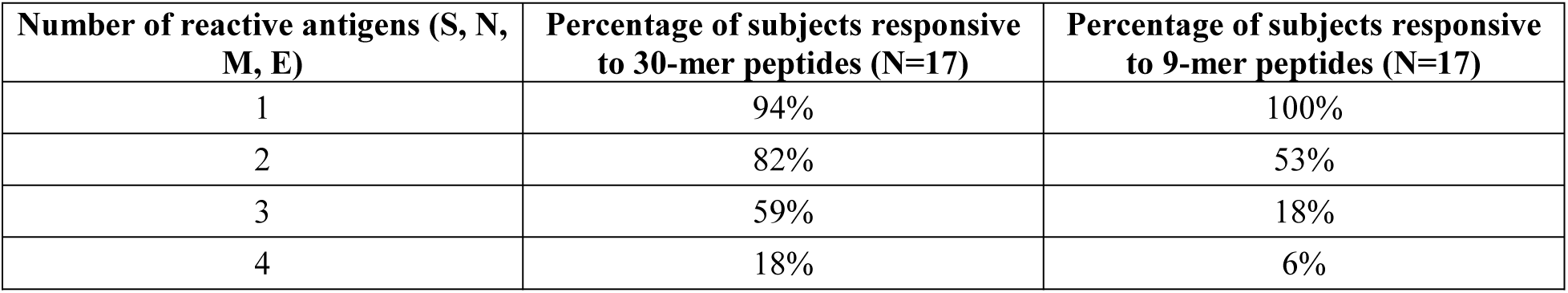
Response rate of COVID-19 convalescent donor patients to one, two, three, or all four viral antigens targeted by the PolyPEPI-SCoV-2 vaccine, as measured by *ex vivo* FluoroSpot assay. Nine-mers are the hotspot HLA class I PEPIs embedded within each 30-mer vaccine peptide coresponding to the four structural proteins: S, Spike; N, Nucleoprotein; M, membrane; E, envelope proteins.

**Supplementary Figure 1.**
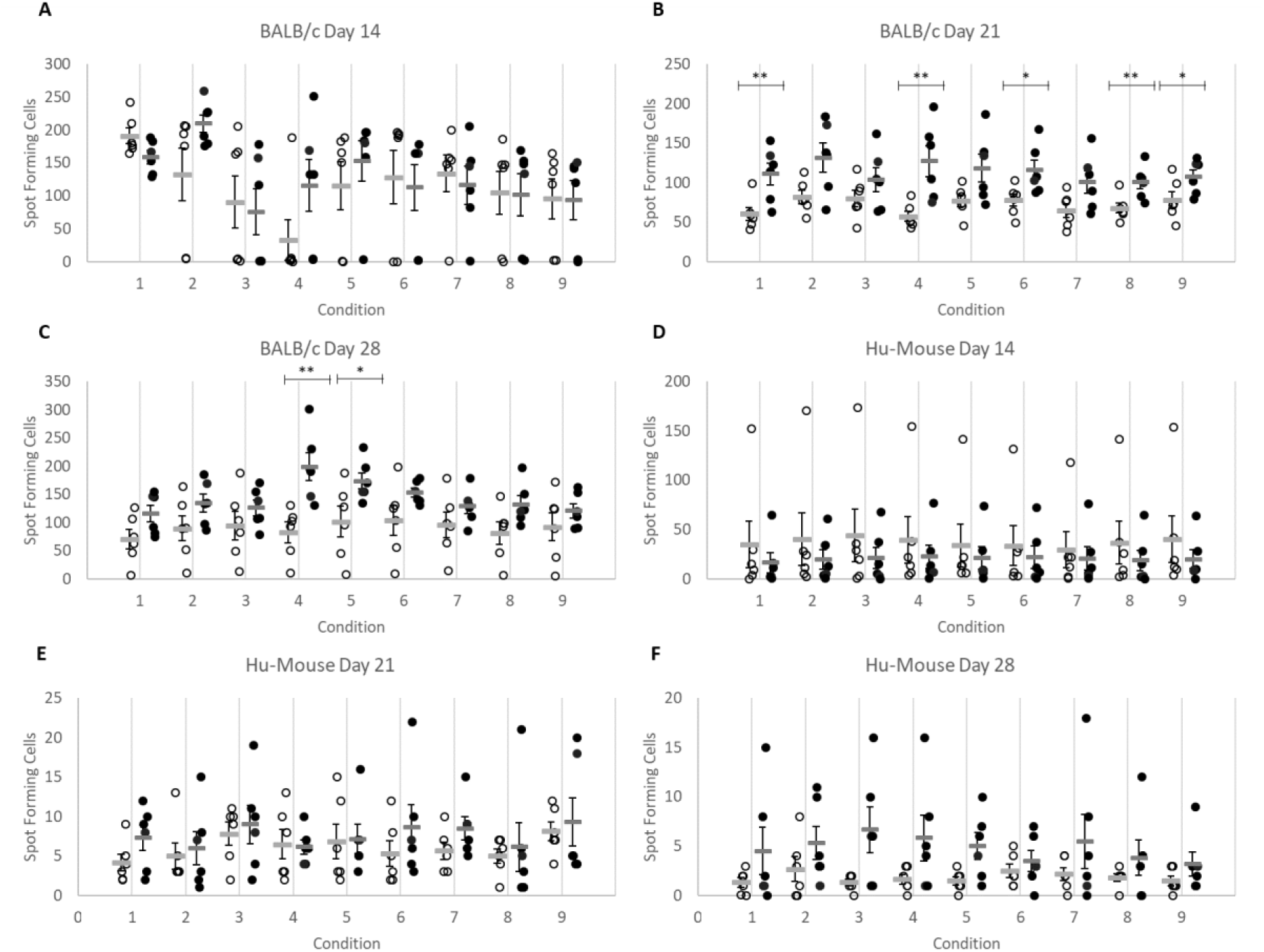
The PolyPEPI-SCoV-2 treatment increases IFN-γ-producing T cells in mice. PolyPEPI-SCoV-2 vaccinated mice are shown with black dots and compared with Vehicle (DMSO/water emulsified with Montanide) control animals shown in white dots. IFN-γ production was analyzed by *ex vivo* ELISpot in the spleen after re-stimulation with peptides at day 14 (**A**, BALB/c; and **D**, Hu-mice), day 21 (**B**, BALB/c; and **E**, Humice), and day 28 (**C**, BALB/c; and **F**, Hu-mice). Condition 1, S-pool; Spike-specific 30-mer pool of S2, S5, and S9 peptides. Condition 2, N-pool; Nucleoprotein-specific 30-mer pool of N1, N2, N3, and N4 peptides. Condition 3, M1 Membrane-specific 30-mer peptide. Condition 4, E1 Envelope-specific 30-mer peptide. Condition 5, S-pool; Spike-specific 9-mer pool of s2, s5, and s9 HLA class I PEPI hotspot fragment of the corresponding 30-mers. Condition 6, N-pool; Nucleoprotein-specific 9-mer pool of n1, n2, n3, and n4 HLA class I PEPI hotspot fragment of the corresponding 30-mers. Condition 7, m1 Membrane-specific 9-mer HLA class I PEPI hotspot fragment of the corresponding 30-mer. Condition 8, e1 Envelope-specific 9-mer HLA class I PEPI hotspot fragment of the corresponding 30-mer. Condition 9, unstimulated control. Individual spot forming cell (SFC) values and means are shown and represent spots per 2^*^10^5^ splenocytes. n=6 mice per group were analyzed. Statistical analysis was performed by Mann-Whitney test. ^*^, p<0.05; ^**^, p<0.01.

**Supplementary Figure 2.**
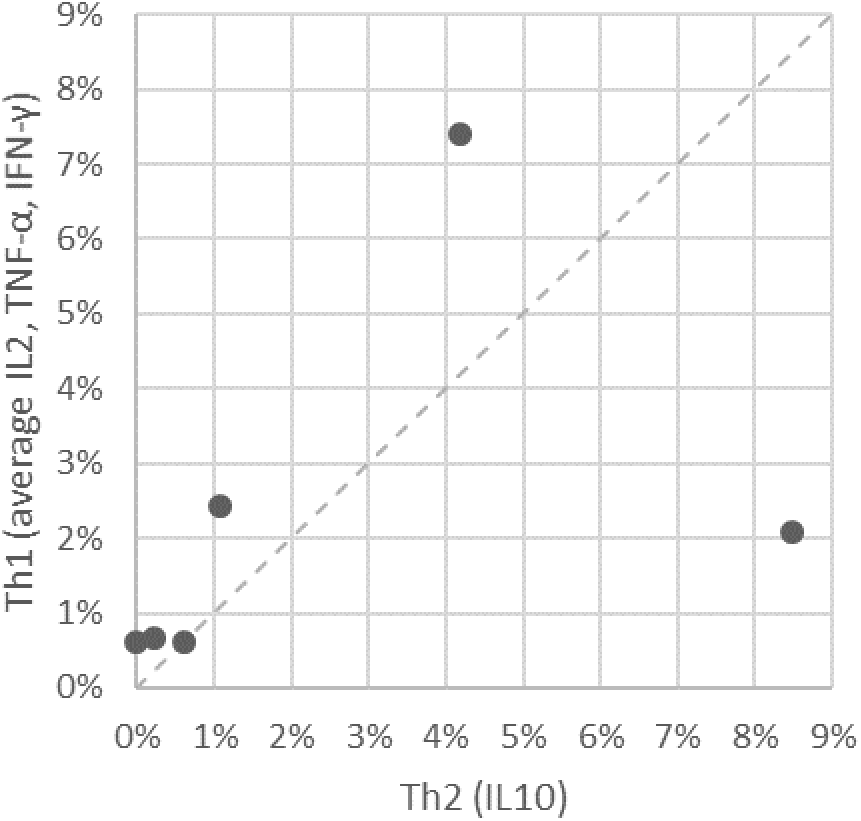
Th1/Th2 balance for T cells detected with PolyPEPI-SCoV-2 vaccine in BALB/c mice at day 28. Average CD4^+^ and CD8^+^ T cells producing IL2, TNF-α, IFN-γ (Th1 cytokines) and IL10 (Th2 cytokine) for each immunized mouse (n=6) using ICS assays. 2 × 10^5^ cells were analyzed, gated for CD45^+^ cells, CD3^+^ T cells, CD4^+^ or CD8^+^ T cells. The average percent was obtained by pooling the background subtracted values of the four stimulation conditions (30-mer S-pool, N-pool, E1 and M1 peptides) for each cytokine for CD4^+^ and CD8^+^ T cells.

**Supplementary Figure 3.**
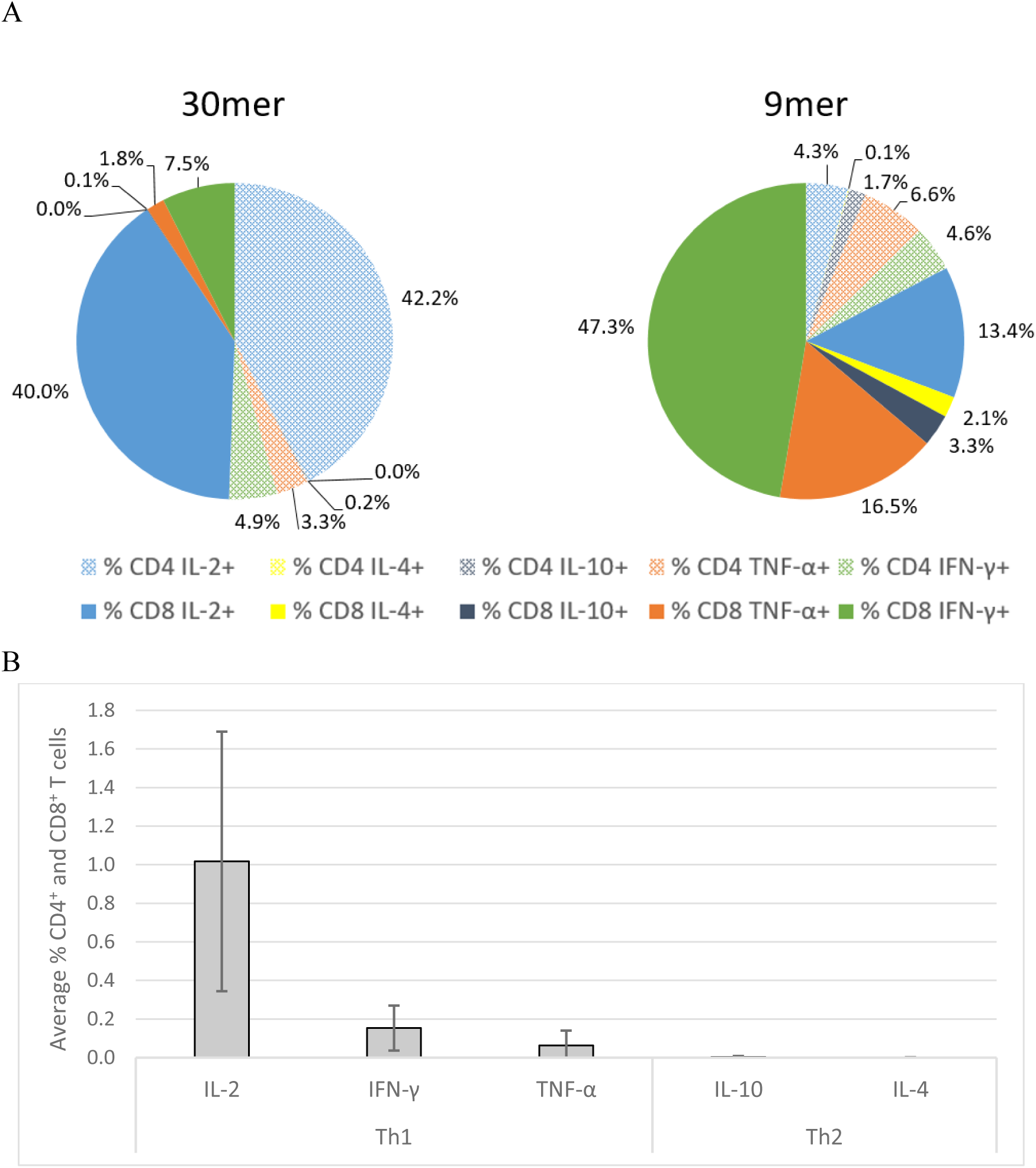
Cytokine production by COVID-19 convalescents’ T cells reactive to PolyPEPI-SCoV-2 peptides determined *ex vivo* from their PBMC by intracellular staining assay. **A)** Cytokine profile of CD4^+^ and CD8^+^ T cells^+^ obtained by stimulations with 9-mer and 30-mer peptides (n=17). **B)** Th1 dominance in vaccine-specific T cells stimulated with 30-mer peptides.

**Supplementary Figure 4.**
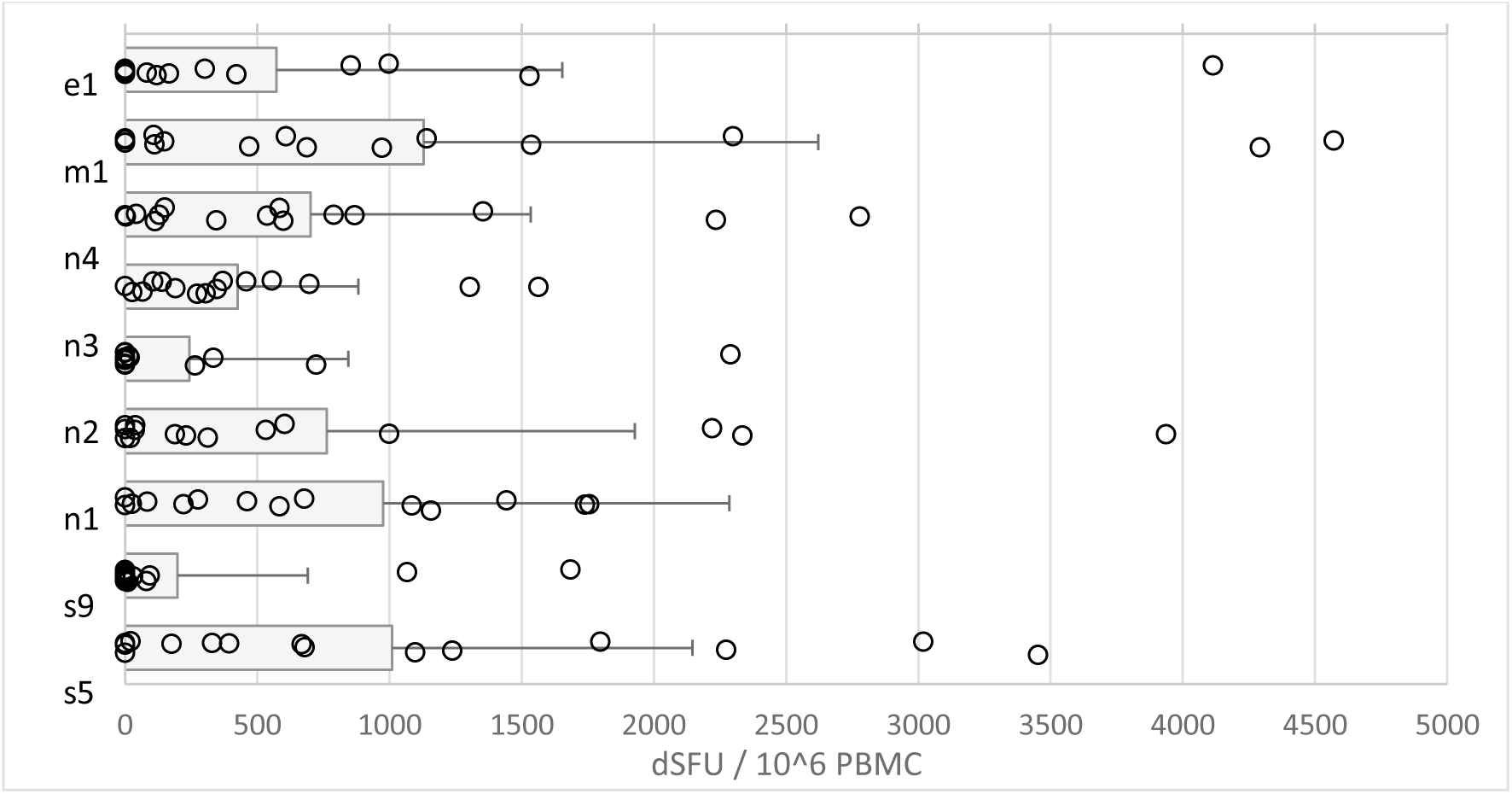
IFN-γ + T cell responses detected for COVID-19 convalescent donors against the 9-mer peptides (PEPI hotspots) of PolyPEPI-SCoV-2 vaccine measured by enriched FluoroSpot assay. S2, S5, and S9 are the three S-specific 9-mer peptide sequences derived from the Spike-specific vaccine 30-mers. N1–N4 are the four Nucleoprotein-specific 9-mer peptide sequences derived from the N-specific vaccine 30-mers. E1 and M1 are Envelope and Membrane-specific 9-mer peptide sequences derived from the E or M-specific vaccine 30-mers, respectively (Table 1 Bold). dSFU, delta spot forming units calculated as non-stimulated background corrected spot counts per 106 PBMC. Average and individual data for each subject are presented. PBMC, peripheral blood mononuclear cells.

**Supplementary Figure 5.**
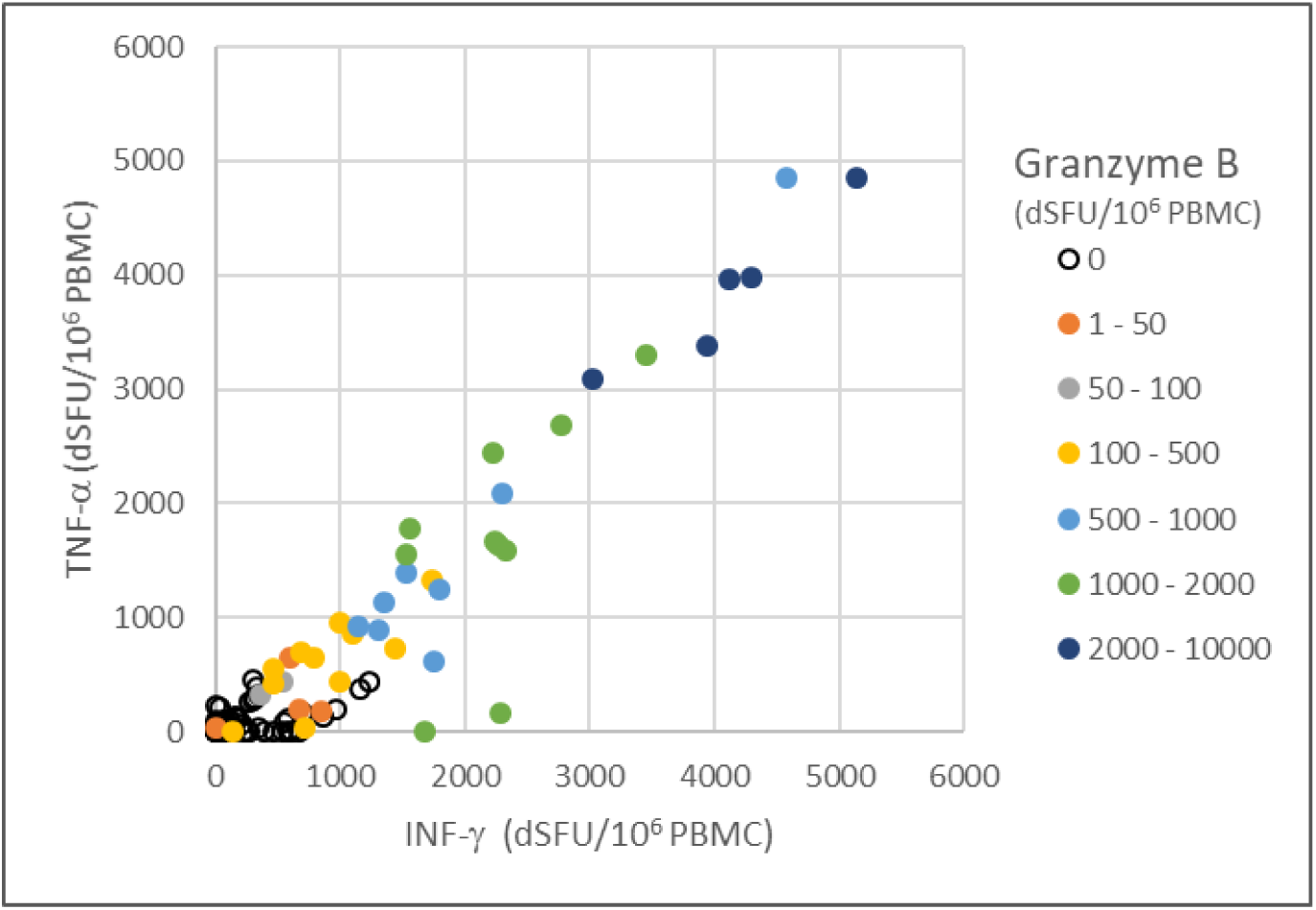
PolyPEPI-SCoV-2-specific polyfunctional T cells detected in COVID-19 convalescents’ blood. IFN-γ and/or TNF-α and/or Granzyme-B positive T cell responses detected for each patient with individual 9-mer peptide stimulations using enriched FluoroSpot assay. dSFU stands for delta spot forming units, calculated as non-stimulated background corrected spot counts per 10^6^ PBMC.

**Supplementary Figure 6.**
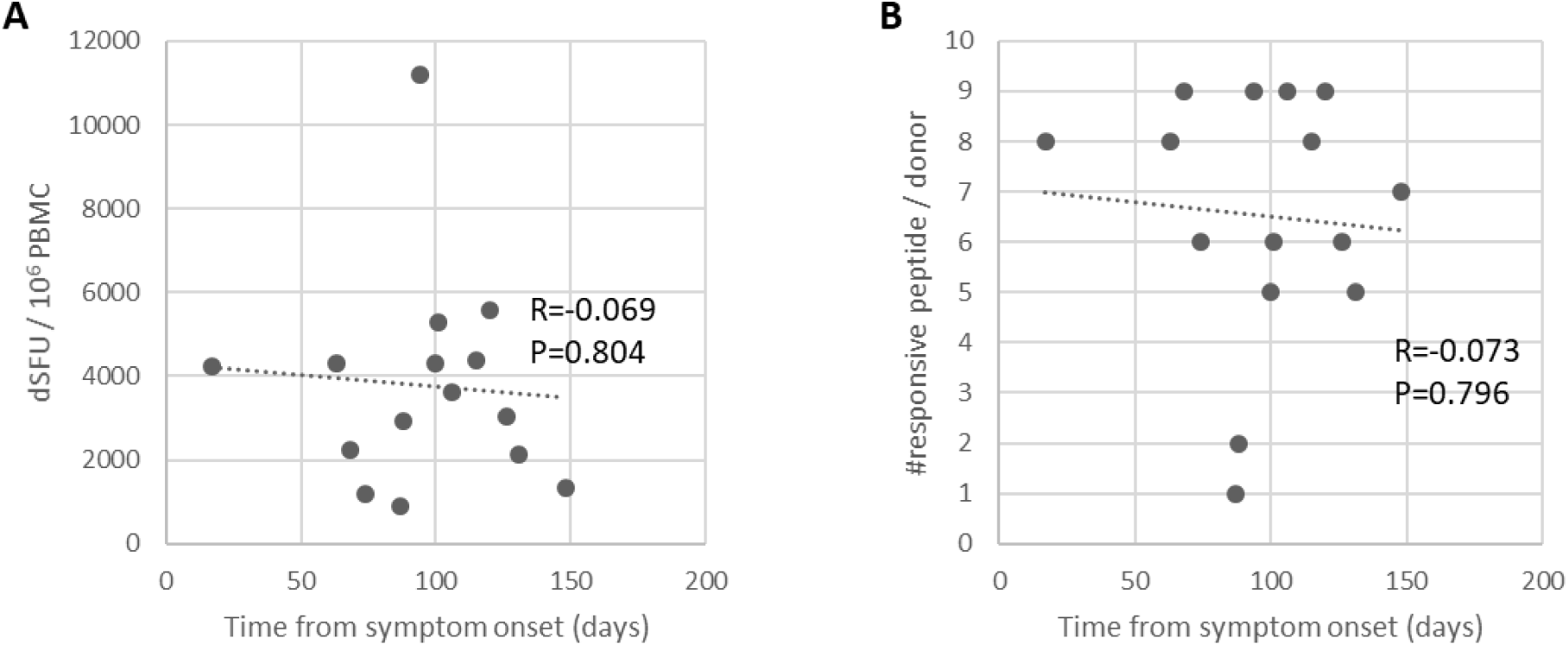
Magnitude and breadth of COVID-19 convalescent donors’T cell responses relative to time from symptom onset. **A)** Magnitude of PolyPEPI-SCoV-2-reactive T cell responses **B)** Breadth of vaccine peptide-reactive CD8^+^ T cell responses from convalescent donors, detected with enriched ELISpot assay. dSFU stands for delta spot forming units, calculated as non-stimulated background corrected spot counts per 10^6^ PBMC. R-Pearson correlation coefficient.

