## Supplementary Tables and Figures for "Peptide vaccine candidate mimics the heterogeneity of natural SARS-CoV-2 immunity in convalescent humans and induces broad T cell responses in mice models"

**Supplementary Table 1. Donor baseline and demographic information.** All donors were caucasoid, with mild/asymptomatic disease and no hospitalization (except one, marked with *). S/Co, sample/control ratio; values were determined according to the manufacturer's instructions, and test results are interpreted as negative in S/Co <0.9, not conclusive if S/CO = 0.9–1.1, and positive if S/Co >1.1. COI, cut-off index; values were determined according to the manufacturer's instructions, and test results are interpreted as negative in COI <0.9, inconclusive with COI 0.9–1.1, and positive if COI >1.1. NA, data not available. Italic, negative or inconclusive values. ** Complaints: a, cough; b, sore throat; c, fever; d, short of breath; e, stomach/intestinal complaints; f, chest pain; g, sore eyes; h, odor or taste loss; i, headache; j, fatigue; k, other complaints (pulmonary embolism and cardiac arrest for IMXP00759; leg pain, arm pain, muscle pain, pain in the eyes).

| Donor ID | Gender | Complaints from/ to: (as reported by the donors) | Complaints** | Blood collection date | Time from first symptom to blood collection | IgA | IgM | IgG | IgG-S1 | IgG-N |
|---|---|---|---|---|---|---|---|---|---|---|
| | | | | | | DiaPro ELISA | | | EUROIMMUN | ROCHE |
| | | | | | | S/Co | S/Co | S/Co | S/Co | COI |
| IMXP00394 | Female | 30 March 2020 - 20 April 2020 | a,b,c,d,h,i,j | 4-Aug-20 | 126 days | *0.36* | 5.065 | 5.752 | 4,48 | 54.38 |
| IMXP00714 | Male | 1 May 2020 – 15 May 2020 | a,b,c,h,i,j,k | 27-Jul-20 | 87 days | 1.324 | 8.524 | 11.524 | 5,35 | 73.06 |
| IMXP00739 | Female | 30 April 2020 | j | 2-Jun-20 | 63 days | *0.929* | 8.841 | 11.967 | 4,54 | 77.61 |
| IMXP00756 | Female | 2 April 2020 - 12 April 2020 | b,c,d,f,i,j | 9-Jun-20 | 68 days | *0.989* | 4.606 | 12.193 | 3,56 | 78.47 |
| IMXP00757 | Female | 29February 2020 - 14 April 2020 | a,b,c,d,e,h,i,j | 9-Jun-20 | 101 days | 1.154 | 5.847 | 8.701 | 7,62 | 29.47 |
| IMXP00758 | Female | 2 April 2020 - 30 arpril 2020 | c,d,h,i,j | 15-Jun-20 | 74 days | 1.356 | 7.757 | 11.774 | 5,79 | 121.9 |
| IMXP00759* | Male | 13 March 2020 - 28 March 2020 | a,c,d,f,h,i,j,k | 15-Jun-20 | 94 days | 6.307 | 10.666 | 13.838 | 9,27 | 87.09 |
| IMXP00762 | Female | 15 March 2020 – 19 March 2020 | b,c,j | 29-Jun-20 | 106 days | 1.251 | 7.314 | 4.46 | 7,25 | 131.5 |
| IMXP00764 | Female | 16 March 2020 – 2 April 2020 | a,b,e,h,i,j,k | 6-Jul-20 | 115 days | 5.161 | 9.739 | 11.677 | 1,32 | 46.59 |
| IMXP00765 | Female | 29 March 2020 - 15 May 2020 | a,d,e,h,i,j,k | 7-Jul-20 | 100 days | *0.565* | 2.948 | 1.54 | 1,32 | 13.4 |
| IMXP00766 | Female | 20 June 2020 - 23 June 2020 | b,c,h,j | 7-Jul-20 | 17 days | *0.771* | 4.648 | 3.973 | 4,14 | 6.25 |
| IMXP00767 | Female | 10 April 2020 - 10 May 2020 | d,e,f,j,k | 7-Jul-20 | 88 days | *0.88* | 5.402 | 3.459 | 2,37 | 52.29 |
| IMXP00771 | Female | 18 March 2020 – 1 April 2020 | a,d,i,j | 28-Jul-20 | 131 days | *0.791* | 7.775 | 8.322 | 4,04 | 119.4 |
| IMXP00772 | Female | 30 March 2020 - 30 April 2020 | g,k | 28-Jul-20 | 120 days | 1.105 | 4.256 | 2.54 | 1,26 | 10.87 |
| IMXP00776 | Female | 9 March 2020 - 14 March 2020 | c,e,i,j,k | 4-Aug-20 | 148 days | *1.012* | 9.196 | 10.887 | 2,26 | 88.64 |
| PTC1 | Male | 15 April 2020 | e | 13-Jul-20 | 89 days | *0.53* | *0.41* | 2.63 | NA | 18.96 |
| PTC2 | Female | 15 April 2020 | e | 13-Jul-20 | 89 days | *0.45* | *0.35* | 1.49 | NA | 26.09 |

**Supplementary Table 2. Sequence alignment results between PolyPEPI-SCoV-2 and other coronavirus strains.** Sequence comparison was made with 8-mer long peptide matching between the aligned protein sequence pairs, defined as the minimum length requirement for a CD8$^+$ T cell epitope. Max AA matching: the longest identical amino acid sequence length. Highlighted grey values represent identical sequences of at least eight amino acids.

| Target Protein | PolyPEPI-SCoV-2 vaccine peptide sequences | | Common 8-mer % / Max AA matching | | | | | |
|---|---|---|---|---|---|---|---|---|
| | | | 229E | NL63 | OC43 | HKU1 | MERS | SARS |
| Spike | S2 | GVYYPDKVFRSSVLHSTQDLFLPFFSNVTW | 0% / 4 | 0% / 4 | 0% / 4 | 0% / 3 | 0% / 3 | 4% / 8 |
| | S5 | DSSSGWTAGAAAYYVGYLQPRTFLLKYNEN | 0% / 3 | 0% / 4 | 0% / 4 | 0% / 4 | 0% / 5 | 0% / 4 |
| | S9 | ALQIPFAMQMAYRFNGIGVTQNVLYENQKL | 0% / 4 | 0% / 4 | 0% / 4 | 0% / 4 | 0% / 5 | 96% / 29 |
| Nucleoprotein | N1 | RSKQRRPQGLPNNTASWFTALTQHGKEDLK | 0% / 3 | 0% / 3 | 0% / 3 | 0% / 3 | 0% / 6 | 78% / 25 |
| | N2 | SKKPRQKRTATKAYNVTQAFGRRGPEQTQG | 0% / 4 | 0% / 4 | 0% / 6 | 0% / 7 | 0% / 4 | 65% / 17 |
| | N3 | ELIRQGTDYKHWPQIAQFAPSASAFFGMSR | 0% / 3 | 0% / 4 | 0% / 3 | 0% / 4 | 0% / 5 | 96% / 29 |
| | N4 | QRQKKQQTVTLLPAADLDDFSKQLQQSMSS | 0% / 5 | 0% / 3 | 0% / 4 | 0% / 3 | 0% / 3 | 9% / 9 |
| Membrane | M1 | LSYFIASFRLFARTRSM**WSFNPETN**ILLNV | 0% / 5 | 0% / 6 | 4% / 8 | 4% / 8 | 4% / 8 | 78% / 25 |
| Envelope | E1 | NIVNVSLVKPSFYVYSRVKNLNSSRVPDLL | 0% / 4 | 0% / 4 | 0% / 3 | 0% / 5 | 0% / 4 | 35% / 12 |

**Supplementary Table 3. Response rate of COVID-19 convalescent donor patients to one, two, three, or all four viral antigens targeted by the PolyPEPI-SCoV-2 vaccine, as measured by *ex vivo* FluoroSpot assay.** Nine-mers are the hotspot HLA class I PEPIs embedded within each 30-mer vaccine peptide coresponding to the four structural proteins: S, Spike; N, Nucleoprotein; M, membrane; E, envelope proteins.

| Number of reactive antigens (S, N, M, E) | Percentage of subjects responsive to 30-mer peptides (N=17) | Percentage of subjects responsive to 9-mer peptides (N=17) |
|---|---|---|
| 1 | 94% | 100% |
| 2 | 82% | 53% |
| 3 | 59% | 18% |
| 4 | 18% | 6% |

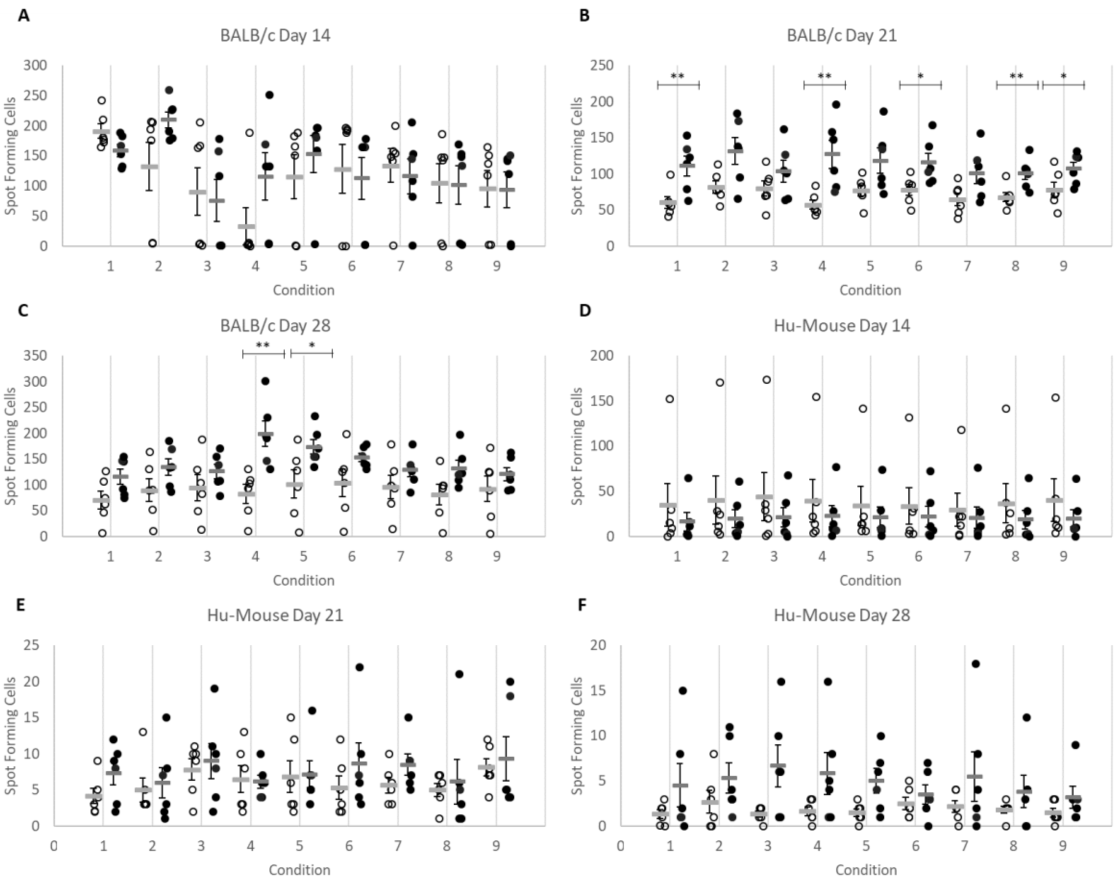


**Supplementary Figure 1. The PolyPEPI-SCoV-2 treatment increases IFN-γ-producing T cells in mice.** PolyPEPI-SCoV-2 vaccinated mice are shown with black dots, and compared to Vehicle (DMSO/Water emulsified with Montanide) control animals shown in white dots. IFN-γ production was analyzed by *ex vivo* ELISpot in the spleen after re-stimulation with peptides at day 14 (**A**, BALB/c; and **D**, Hu-mice), day 21 (**B**, BALB/c; and **E**, Hu-mice), and day 28 (**C**, BALB/c; and **F**, Hu-mice). Condition 1, S-pool; Spike-specific 30-mer pool of S2, S5, and S9 peptides. Condition 2, N-pool; Nucleoprotein-specific 30-mer pool of N1, N2, N3, and N4 peptides. Condition 3, M1 Membrane-specific 30-mer peptide. Condition 4, E1 Envelope-specific 30-mer peptide. Condition 5, S-pool; Spike-specific 9-mer pool of s2, s5, and s9 HLA class I PEPI hotspot fragment of the corresponding 30-mers. Condition 6, N-pool; Nucleoprotein-specific 9-mer pool of n1, n2, n3, and n4 HLA class I PEPI hotspot fragment of the corresponding 30-mers. Condition 7, m1 Membrane-specific 9-mer HLA class I PEPI hotspot fragment of the corresponding 30-mer. Condition 8, e1 Envelope-specific 9-mer HLA class I PEPI hotspot fragment of the corresponding 30-mer. Condition 9, unstimulated control. Individual spot forming cell (SFC) values and means are shown and represent spots per $2*10^5$ splenocytes. n=6 mice per group were analyzed. Statistical analysis was performed by Mann-Whitney test. *, $p<0.05$; **, $p<0.01$.

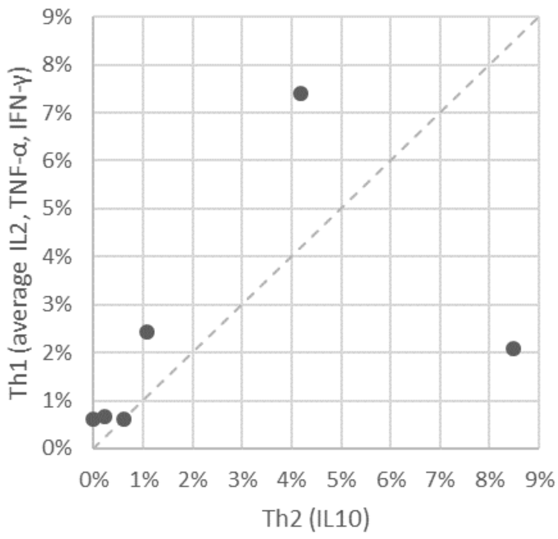


**Supplementary Figure 2. Th1/Th2 balance for T cells detected with PolyPEPI-SCoV-2 vaccine in BALB/c mice at day 28.** Average $CD4^+$ and $CD8^+$ T cells producing IL2, TNF-α, IFN-γ (Th1 cytokines) and IL10 (Th2 cytokine) for each immunized mice (n=6) using ICS assay. $2*10^5$ cells were analyzed, gated for $CD45^+$ cells, $CD3^+$ T cells, $CD4^+$ or $CD8^+$ T cells. The average percent was obtained by pooling the background subtracted values of the 4 stimulation conditions (30-mer S-pool, N-pool, E1 and M1 peptides) for each cytokine for $CD4^+$ and $CD8^+$ T cells

A

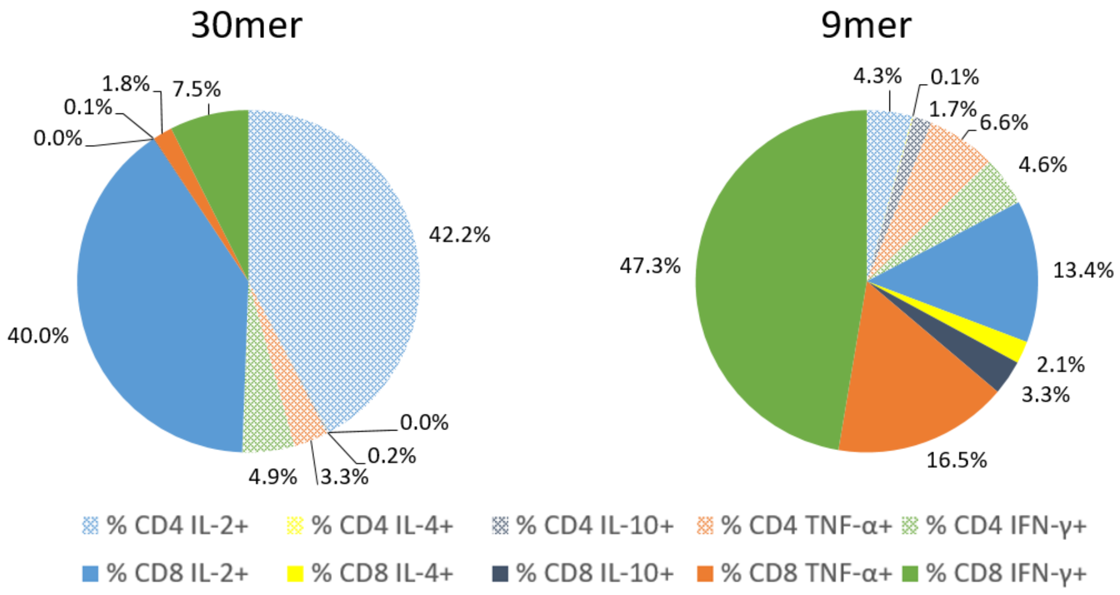


B

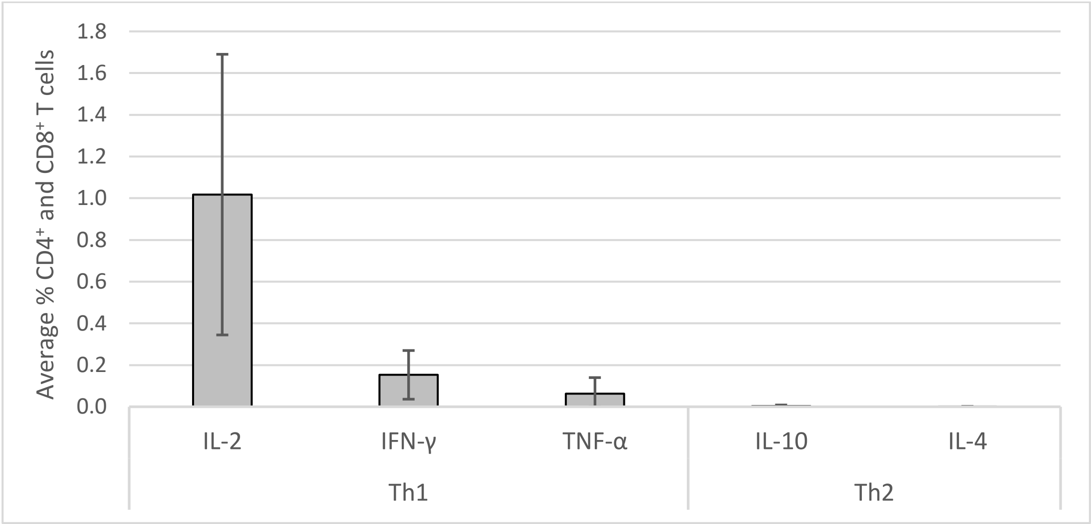


**Supplementary Figure 3. Cytokine production by COVID-19 convalescents' T cells reactive to PolyPEPI-SCoV-2 peptides determined *ex vivo* from their PBMC by intracellular staining assay. A)** Cytokine profile of $CD4^+$ and $CD8^+$ T cells+ obtained by stimulations with 9-mer and 30-mer peptides (n=17). **B)** Th1 dominance in vaccine-specific T cells stimulated with 30-mer peptides.

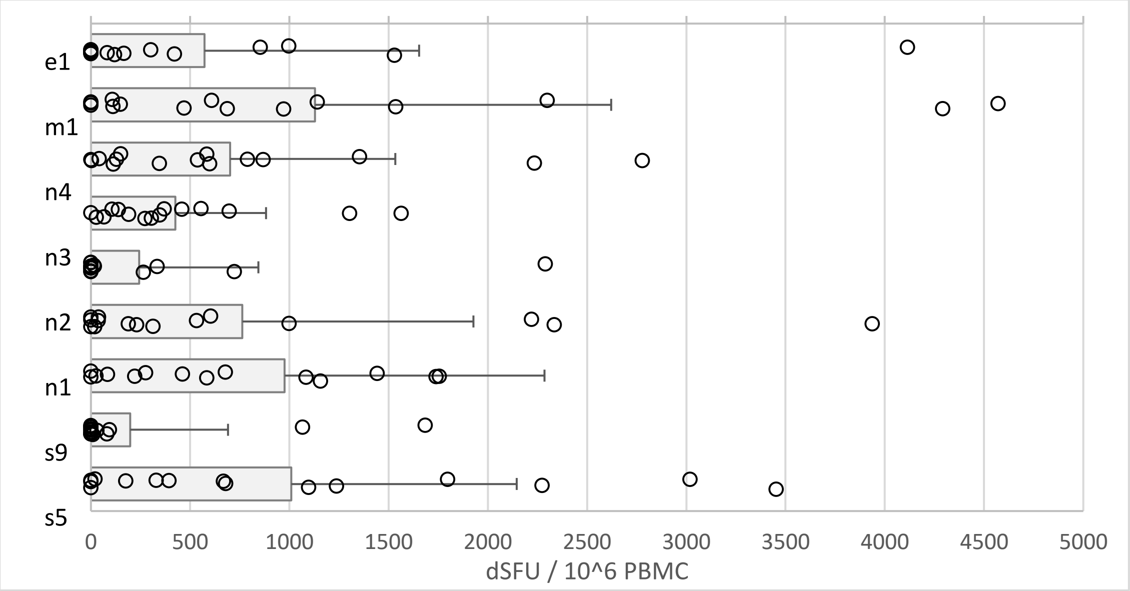


**Supplementary Figure 4. IFN-γ + T cell responses detected for COVID-19 convalescent donors against the 9-mer peptides (PEPI hotspots) of PolyPEPI-SCoV-2 vaccine measured by enriched FluoroSpot assay.** s2, s5, and s9 are the three S-specific 9-mer peptide sequences derived from the Spike-specific vaccine 30-mers. n1–n4 are the four Nucleoprotein-specific 9-mer peptide sequences derived from the N-specific vaccine 30-mers. e1 and m1 are Envelope and Membrane-specific 9-mer peptide sequences derived from the E or M-specific vaccine 30-mers, respectively (Table 1 Bold). dSFU, delta spot forming units calculated as non-stimulated background corrected spot counts per $10^6$ PBMC. Average and individual data for each subject are presented. PBMC, peripheral blood mononuclear cells.

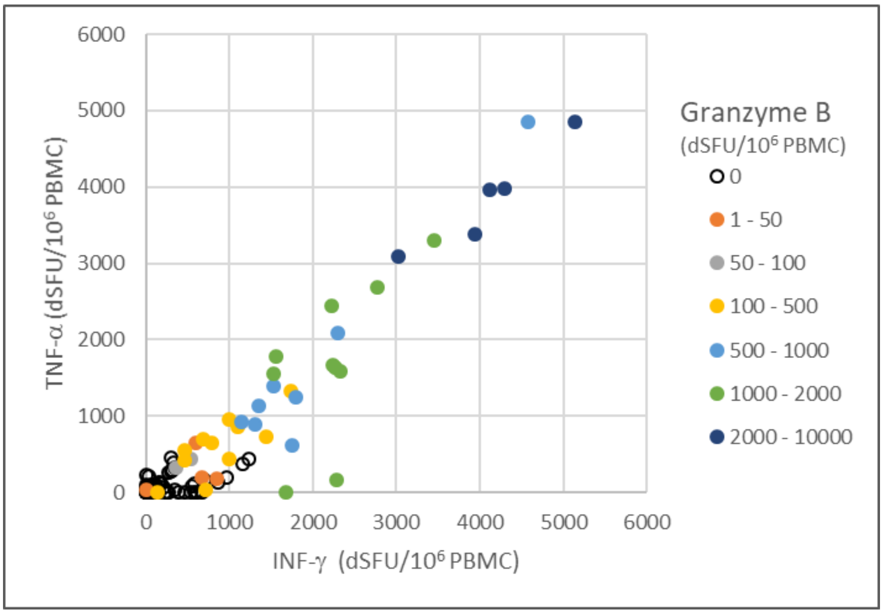


**Supplementary Figure 5. PolyPEPI-SCoV-2-specific polyfunctional T cells detected in COVID-19 convalescents' blood.** IFN-γ and/or TNF-α and/or Granzyme-B positive T cell responses detected for each patient with individual 9-mer peptide stimulations using enriched FluoroSpot assay. dSFU stands for delta spot forming units, calculated as non-stimulated background corrected spot counts per $10^6$ PBMC.

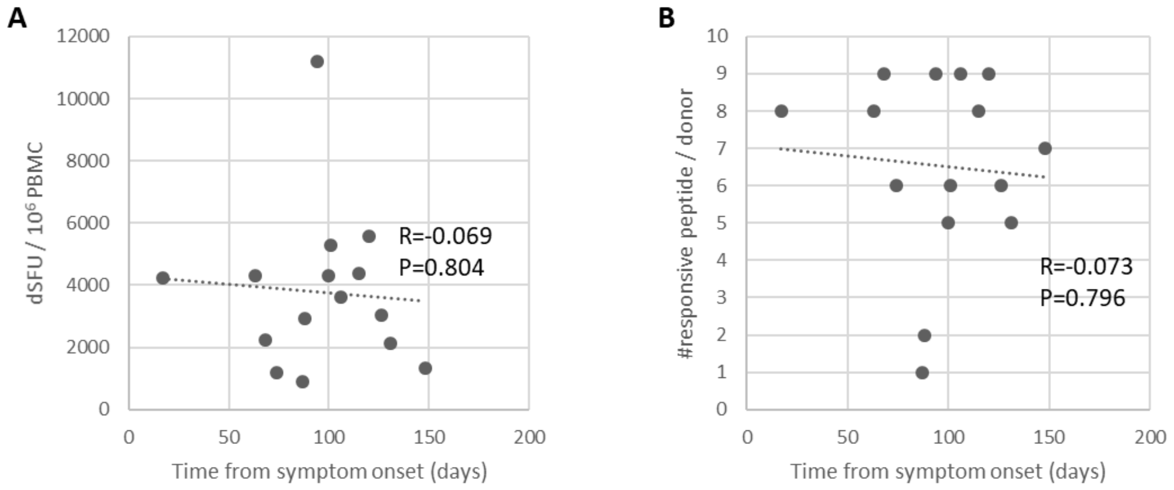


**Supplementary Figure 6. Magnitude and breadth of COVID-19 convalescent donors'T cell responses relative to time from symptom onset. A)** Magnitude of PolyPEPI-SCoV-2-reactive T cell responses **B)** Breadth of vaccine peptide- reactive $CD8^+$ T cell responses from convalescent donors, detected with enriched ELISpot assay. dSFU stands for delta spot forming units, calculated as non-stimulated background corrected spot counts per $10^6$ PBMC. R- Pearson correlation coefficient.
